# EBF1 limits the numbers of cochlear hair and supporting cells and forms the scala tympani and spiral limbus during inner ear development

**DOI:** 10.1101/2023.04.28.538789

**Authors:** Hiroki Kagoshima, Hiroe Ohnishi, Ryosuke Yamamoto, Akiyoshi Yasumoto, Yosuke Tona, Takayuki Nakagawa, Koichi Omori, Norio Yamamoto

## Abstract

Early B-cell factor 1 (EBF1) is a basic helix-loop-helix transcription factor essential for the differentiation of various tissues. Our single-cell RNA sequencing data suggest that *Ebf1* is expressed in the sensory epithelium of the inner ear. Here, we found that the *Ebf1* gene and its protein are expressed in the prosensory domain of the inner ear, medial region of the cochlear floor, inner ear mesenchyme, and cochleo-vestibular ganglion. *Ebf1* deletion results in incomplete formation of the spiral limbus and scala tympani, increased number of cells in the organ of Corti and Kölliker’s organ, and aberrant course of the spiral ganglion axons. The inner ear-specific deletion of *Ebf1* revealed that scala tympani formation depended on *Ebf1* expressed in the otic mesenchyme. *Ebf1* deletion in the cochlear epithelia caused the proliferation of SOX2-positive cochlear cells at E13.5, indicating that EBF1 suppresses the proliferation of the prosensory domain and cells of Kölliker’s organ to facilitate the development of appropriate numbers of hair and supporting cells. Furthermore, mice with deletion of cochlear epithelium-specific *Ebf1* showed poor postnatal hearing function. Our results suggest that *Ebf1* is essential for normal auditory function in mammals.

## Introduction

The inner ear is a unique and complex organ that consists of bony and membranous labyrinths. The membranous labyrinth contains multiple sensory organs, including the cochlea, which is responsible for hearing and comprises three compartments (scalae): the scala vestibuli, scala tympani, and scala media. The scala vestibuli and scala tympani develop from the mesenchyme surrounding the inner ear (Sher, 1971). The scala media is situated between the scala vestibuli and scala tympani, which are separated from the scala media by the tectorial and basilar membranes, respectively. Moreover, the scala media contains sensory epithelia that transduce sound into electrical signals via specialized sensory cells known as hair cells. Cochlear hair cells are located within the organ of Corti in the middle part of the scala media epithelium, and consist of one row of inner hair cells and three rows of outer hair cells. These hair cells are surrounded by at least seven types of non-sensory supporting cells: inner boundary cells, inner phalangeal cells, Deiters’ cells, inner and outer pillar cells, Hensen’s cells, and Claudius cells. The precise number and placement of mechanosensory hair cells and non-sensory supporting cells enable the accurate reception of mechanical stimulation of sound or body motion and its conversion into neural signals.

Inner ear development in mice begins with the formation of an ectodermal thickening called the otic placode, which is located adjacent to the hindbrain (Wu and Kelley, 2012). The otic placode invaginates to form a spherical structure called an otocyst at approximately embryonic day (E) 9.5. The ventral side of the otocyst forms the future sensory epithelium, where the sex-determining region Y-box transcription factor 2 (*Sox2*) is expressed (Kiernan et al., 2005). At E10.5, the cochlear and endolymphatic ducts and semicircular canals begin to form on the ventral and dorsal sides of the otocyst, respectively. After ventral cochlear duct formation becomes evident at E11, the ventral side of the cochlear duct (cochlear duct floor) begins to develop into a future sensory domain by expressing *Sox2* and *Jagged1* at E11.5 (Wu and Kelley, 2012). The *Sox2*-positive region becomes limited to the middle part of the ventral cochlear duct and is recognized as a prosensory domain at E13.5 and E14.5 (Kiernan et al., 2005; Ohyama et al., 2010). This region is located between two molecularly distinct nonsensory areas. The region medial to the prosensory domain toward the axis of the cochlea (modiolus) is called the greater epithelial ridge (GER) and transiently contains Kölliker’s organ, which is composed of columnar supporting cells during the developmental stage and becomes the inner sulcus with cuboidal cells and the spiral limbus with interdental cells in the mature cochlea (Dayaratne et al., 2014). Additionally, the GER is a source of sensory epithelia and has the potential to produce sensory cells after the establishment of hair cells (Kubota et al., 2021).

The complex cellular structure and developmental processes of the inner ear depend on the highly regulated expression patterns of signaling molecules and transcription factors. However, the mechanisms underlying inner ear development are not fully understood. To comprehensively elucidate these mechanisms, we analyzed the single-cell RNA-seq data of the inner ear epithelial cells. In this study, we found that the early B-cell factor 1 gene (*Ebf1*) was upregulated in clusters of sensory epithelial progenitors in the cochlea, macula, and crista, and confirmed that it was expressed on the medial side of the cochlear duct floor, the prosensory area of the macula and crista, and the spiral ganglion (Yamamoto et al., 2021).

EBF1 belongs to the EBF family of transcription factors and encodes four paralogous genes in mammals, including mice and humans (Liberg et al., 2002). EBF family proteins comprise a functional DNA-binding domain, transcription factor immunoglobulin, helix-loop-helix (HLH) domain, and transactivation domain (Hagman et al., 1995), which are the characteristics of basic HLH (bHLH) transcription factors. Similar to other bHLH transcription factors, EBF1 is involved in various developmental processes, including the determination of cell fate and differentiation of B lymphocytes and olfactory epithelia (Liberg et al., 2002).

Considering the various roles of EBF1 as a bHLH transcription factor, the importance of bHLH transcription factors—such as ATOH1—in inner ear development, and the results of our previous study confirming *Ebf1* expression in the inner ear sensory epithelium, we analyzed the function of EBF1 in inner ear development. In the present study, we confirmed the spatiotemporal expression of *Ebf1* during inner ear development and examined the effects of *Ebf1* deletion on inner ear development and hearing.

## Results

### Ebf1 is expressed in developing mouse cochlea

*In silico* analysis of embryonic inner ear epithelia suggested that *Ebf1* is predominantly expressed in the inner ear sensory epithelium during early development (Yamamoto et al., 2021). To quantify *Ebf1* expression at each stage of inner ear development, we performed quantitative reverse transcription polymerase chain reaction (qRT-PCR) using whole embryonic inner ears from E9.5 to P0 (Fig. 1A). The expression of *Ebf1* mRNA transcripts began to increase at E10.5 and reached a maximum at E13.5 (Fig. 1A). The relative expression level at E13.5 was approximately 10-fold higher than that at E9.5. The expression level then decreased but remained 7.5 times higher than that at E9.5 even at P0.

**Fig. 1.**
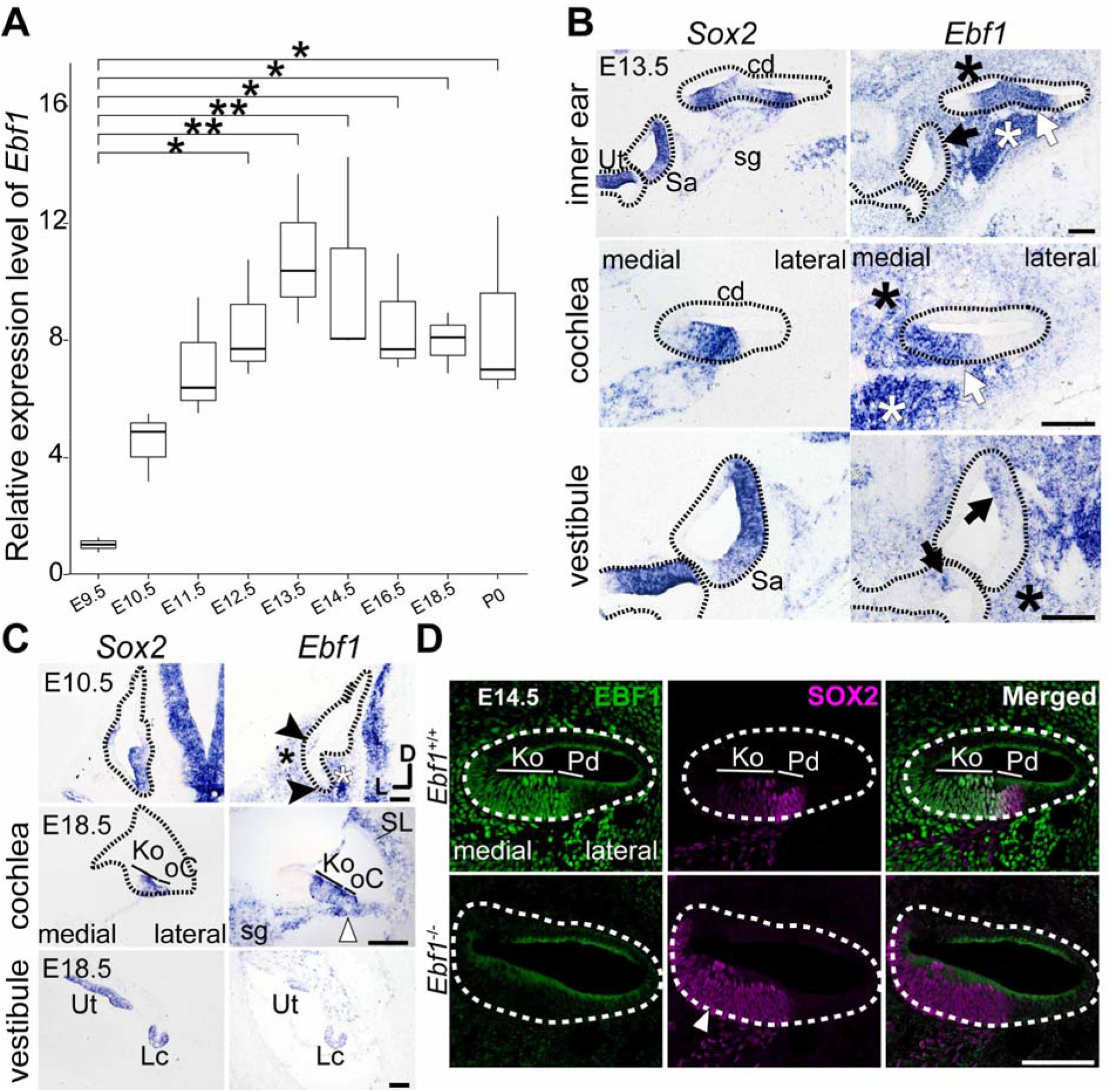
Quantitative and spatiotemporal expression of *Ebf1* during inner ear development. **A.** Results of quantitative reverse transcription polymerase chain reaction (qRT-PCR) analysis for *Ebf1* in the inner ear of wild-type mice from embryonic day (E) 9.5 to P0. The value of each date is normalized to the value of E9.5. Box plot representing the medians and interquartile ranges of the relative mRNA expression of *Ebf1*. **B** and **C**. Result of *in situ* hybridization for *Ebf1* and *Sox2* in cross sections of the inner ear of wild-type mice at E13.5 (B) and E10.5 and E18.5 (C). Areas enclosed by dashed lines indicate the inner ear epithelium. Low-magnification images of the cochlear basal turn and vestibule are presented in the uppermost panels of **B**. High-magnification images of the apical turn of the cochlear duct and vestibules are presented in the middle and lower panels of **B**. Images of E10.5 otocysts and E18.5 cochleae and vestibules are presented in the uppermost, middle, and lowermost images of **C**, respectively. **D**. Immunohistochemical images of E14.5 *Ebf1^+/+^* (upper panels) and *Ebf1^-/-^* (lower panels) mouse cochlear ducts. One-way analysis of variance (ANOVA) with Tukey–Kramer *post-hoc* tests was performed. \**p* <0.05, \*\**p* <0.01. D, dorsal; L, lateral. cd, cochlear duct; sg, spiral ganglion; Ut, utricle; Sa, saccule; Lc, lateral crista; SL, spiral ligament; Ko, Kölliker’s organ; oC, organ of Corti; Pd, prosensory domain. Scale bars: 100 μm.

To describe the spatiotemporal expression patterns of *Ebf1* during inner ear development, we performed *in situ* hybridization (ISH) and immunohistochemistry (IHC) analyses on sections of the developing inner ear of wild-type mice from E9.5 to P0 (Fig. 1 B–D, Supplemental Fig. S1). We stained *Sox2*, which is expressed in the sensory progenitor region of the inner ear from the early developmental stages (Kiernan et al., 2005), as well as *Ebf1* on adjacent sections to specify the location of *Ebf1* expression, and compared the expression of the two genes and their products. First, we examined *Ebf1* expression at E13.5 (Fig. 1B), which is when the sensory epithelium of the inner ear forms and *Ebf1* expression level is maximized during inner ear development (Fig. 1A). *Ebf1* was expressed on the medial side of the cochlear floor, including the prosensory domain (white arrows in Fig. 1B), spiral ganglion (white asterisks in Fig. 1B), otic mesenchyme (black asterisks in Fig. 1B), and parts of the prosensory regions of the vestibule and crista (black arrows in Fig. 1B). Compared with *Sox2*, *Ebf1* was expressed more medially within the cochlear floor, which developed into Kölliker’s organ and the organ of Corti, and its expression in the vestibule was more restricted (Fig. 1B).

Subsequently, we examined the spatiotemporal expression of *Ebf1* throughout inner ear development, including the onset of expression in the inner ear epithelium, using inner ear sections from E9.5 to P0 (Fig. 1C, Supplemental Fig. S1). At E9.5, *Ebf1* was not expressed in the otocyst but was expressed in the progenitor cells of the cochleo-vestibular ganglion (CVG) (data not shown), which delaminate from the ventral side of the otocyst into the otic mesenchyme (Wu and Kelley, 2012). At E10.5, *Ebf1* expression was observed on the ventromedial side of the otocyst, which develops into the cochlear duct, and in the ventrolateral epithelium of the otocyst, which develops into the crista (black arrowheads in Fig. 1C). Additionally, *Ebf1* expression was detected in the otic mesenchyme (black asterisk in Fig. 1C) and CVG (white asterisk in Fig. 1C), which persisted until later stages (Supplemental Fig. S1). *Ebf1* was expressed in the border region, where the cochlear duct begins to elongate, at E11.5, in the medial side of the cochlear floor at E12.5, and in the future crista region in the vestibule at E11.5 and 12.5 (Supplemental Fig. S1). In the cochlea at E16.5 and E18.5, *Ebf1* was expressed throughout the organ of Corti and Kölliker’s organ, whereas *Sox2* was expressed in the organ of Corti and the lateral half of Kölliker’s organ (Fig. 1C and Supplemental Fig. S1), consistent with a previous report (Urness et al., 2015). *Ebf1* was expressed in the spiral ligaments, tympanic border cells (white arrowheads in Fig. 1C) (Taniguchi et al., 2012), vestibules, and crista (Fig. 1C). *Ebf1* expression was maintained until P0 (Supplemental Fig. S1).

IHC analysis showed that EBF1 was expressed throughout Kölliker’s organ and the prosensory domain, whereas SOX2 was expressed in a part of Kölliker’s organ and the prosensory domain (upper panels of Fig. 1D), which is consistent with the ISH results. The disappearance of the EBF1 signal from the cochlear epithelia and mesenchyme in conventional *Ebf1* knockout (*Ebf1^-/-^*) mice confirmed the specificity of the anti-EBF1 antibody used in this study (Fig. 1D).

### Ebf1 deletion altered the structure of the cochlear duct

Our ISH and IHC analyses, which showed the expression of *Ebf1* and its protein in both the developing inner ear epithelia and mesenchyme, suggest that *Ebf1* is involved in the development of both the inner ear sensory epithelium and otic mesenchyme. We used two mutant mouse strains to examine the roles of *Ebf1* in developing inner ears: an *Ebf1* conventional knockout (*Ebf1^-/-^*) mouse (Lin and Grosschedl, 1995) and a Foxg1-Cre-mediated inner ear epithelia-specific conditional knockout mouse (*Foxg1Cre;Ebf1^fl/fl^*) (Hébert and McConnell, 2000; Gyory et al., 2012) in which *Ebf1* expression remains localized in the inner ear mesenchyme.

Comparison of the gross morphology of the membranous labyrinth of the inner ear at E18.5 revealed no difference between *Ebf1^+/+^* and *Ebf1^+/-^* mice (Fig. 2A). However, after removing the lateral wall and roof of the cochlea to expose the cochlear floor, we found that *Ebf1^-/-^*mice had fewer turns in the cochlear duct than *Ebf1^+/+^* mice (white arrowhead in Fig. 2B). Hematoxylin-eosin (HE) staining of the cochlea at E18.5 revealed incomplete formation of the scala tympani, particularly in the middle and apical turns of the cochlea of *Ebf1*^-/-^ mice compared with those of control mice (arrowheads in Fig. 2C). Moreover, *Ebf1*^-/-^ mice lacked a spiral limbus (sl; Fig. 2C); a lower number of cochlear turns was also observed in the cochlear sections of these mice (arrows in the upper right panels in Fig. 2C), supporting the gross morphological observations.

**Fig. 2.**
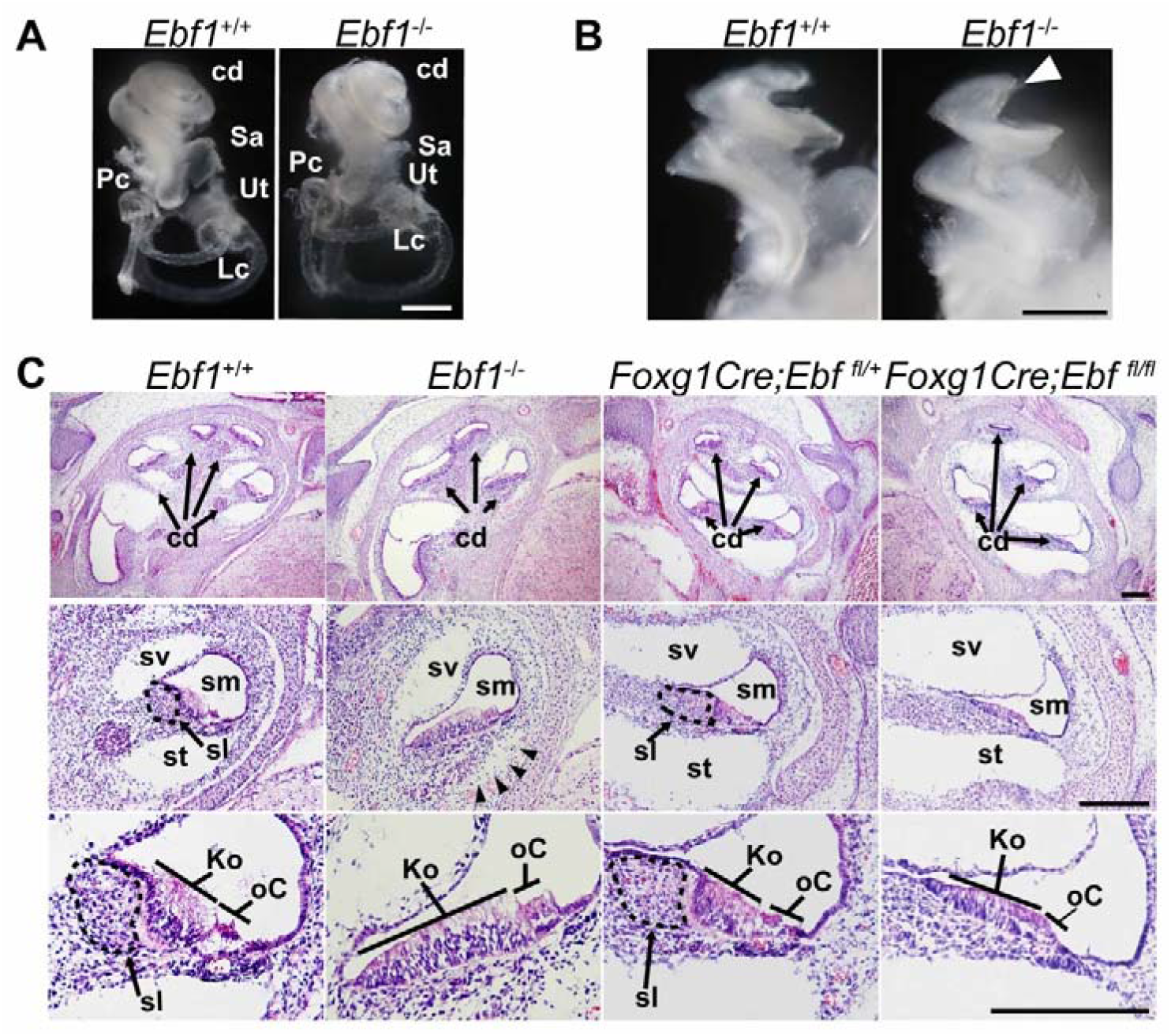
Development of the cochlear duct is deteriorated in *Ebf1^-/-^* mice. **A.** Gross morphology of membranous labyrinth of the inner ear of *Ebf1^+/+^*and *Ebf1*^−/−^mice at embryonic day (E) 18.5. **B.** Gross morphology of the cochlear floor after the lateral wall and roof of the cochlea were removed at E18.5. **C.** Hematoxylin and eosin-stained cross sections of the cochlea at E18.5 from *Ebf1^+/+^*, *Ebf1*^−/−^, *Foxg1Cre;Ebf1^fl/+^*, and *Foxg1Cre;Ebf1^fl/fl^* mice. The upper panels are low-magnification images, and the middle and lower panels are high-magnification images. Areas enclosed by dashed lines indicate the spiral limbus. cd, cochlear duct; Ut, utricle; Sa, saccule; Lc, lateral crista; Pc, posterior crista; sv, scala vestibuli; sm, scala media; st, scala tympani; sl, spiral limbus: Ko, Kölliker’s organ; oC, organ of Corti. Scale bars: 0.5 mm in **A** and **B** and 200 μm in **C**.

*Ebf1* was expressed in the otic mesenchyme from early to later developmental stages. To elucidate whether the hypoplastic scala tympani, fewer cochlear turns, and lack of spiral limbus were caused by the *Ebf1*-deficient mesenchyme, we examined the cochlear morphology of *Foxg1Cre;Ebf1^fl/fl^* and *Foxg1Cre;Ebf1^fl/+^*mice via HE staining. In contrast to the hypoplastic scala tympani of *Ebf1*^-/-^ mice, the scala tympani of *Foxg1Cre;Ebf1^fl/fl^* mice was completely formed throughout the cochlear turns (left panels of Fig. 2C). The number of cochlear turns in *Foxg1Cre;Ebf1^fl/fl^* mice was similar to that in the control mice (*Foxg1Cre;Ebf1^fl/+^* mice). However, *Foxg1Cre;Ebf1^fl/fl^* mice lacked a spiral limbus, as observed in *Ebf1*^-/-^ mice (Fig. 2C). These results suggest that *Ebf1* controls scala tympani formation and the turning of the cochlear duct through the mesenchyme, and the lack of the spiral limbus in *Ebf1*-deficient mice is secondarily caused by EBF1 malfunction within the epithelia.

### Ebf1 deletion caused an increase in the number of cochlear hair, supporting, and Kölliker’s organ cells

Observation of the cochlear epithelia in HE-stained samples revealed that both *Ebf1*^-/-^ and *Foxg1Cre;Ebf1^fl/fl^*mice had deformed Kölliker’s organs and organs of Corti (Ko and oC in Fig. 2C). To examine these phenotypes more comprehensively, we performed IHC analysis on inner ear sections and cochlear whole-mount samples from E18.5 (Fig. 3 and Supplemental Figs. S2 and S3).

**Fig. 3.**
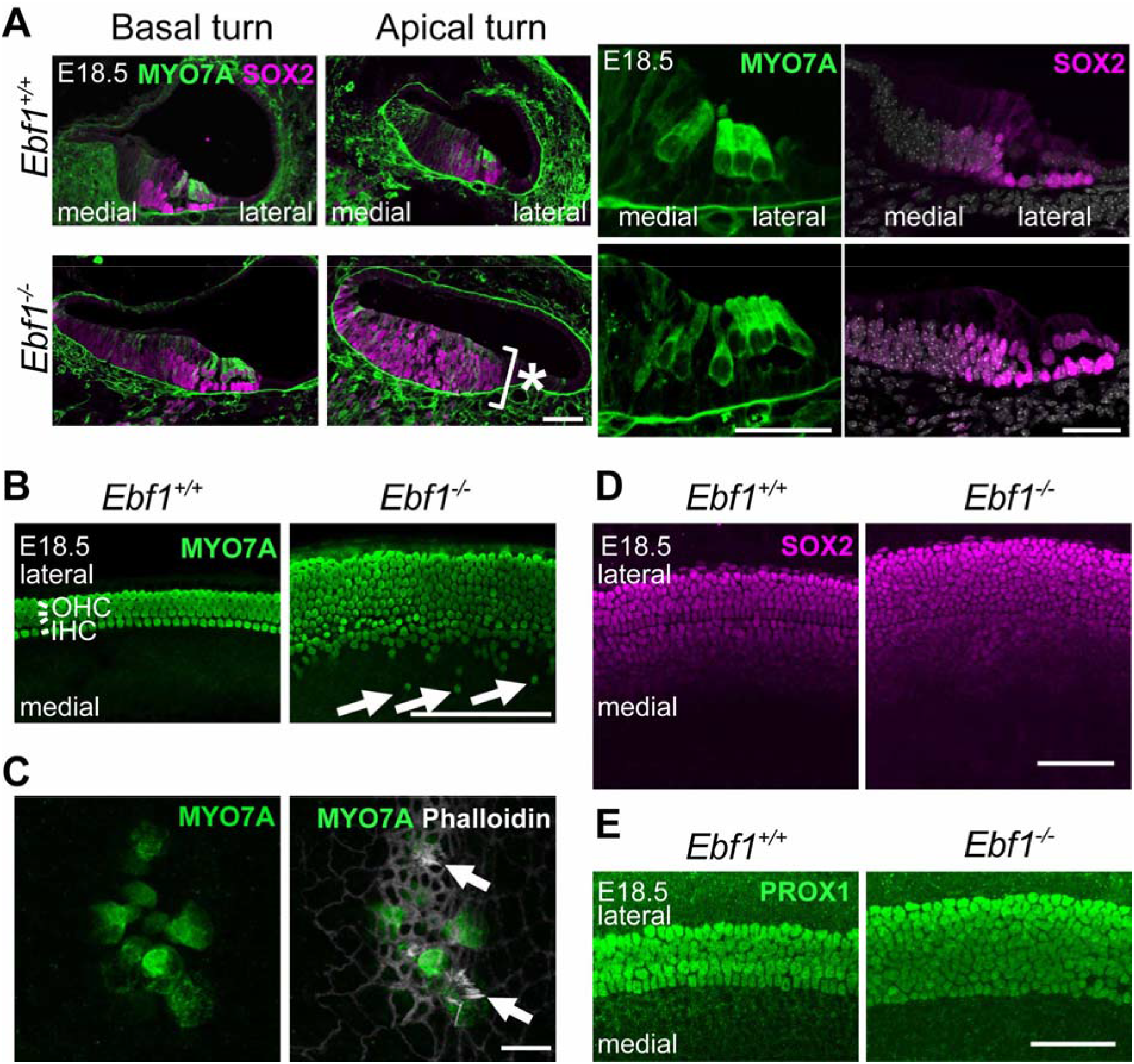
*Ebf1* deletion increases the number of MYO7A-, SOX2-, and PROX1-positive cells in the cochlea. **A.** Cross sections of the basal and apical turns of the cochlea of *Ebf1^+/+^* and *Ebf1^-/-^* mice at embryonic day (E) 18.5, labeled with MYO7A (green) and SOX2 (magenta). Magnified images of the organ of Corti (oC) and Kölliker’s organ (Ko) are presented in the four right panels. **B**–**E**. Whole mount cochlear images from *Ebf1^+/+^* and *Ebf1^-/-^* mice at E18.5, labeled with MYO7A (green, **B** and **C**), phalloidin (white, **C**), SOX2 (magenta, **D**), and PROX1 (green, **E**). Scale bars: 50 μm in **A**, **D**, and **E**; 100 μm in **B**; 10 μm in **C**.

IHC analysis of cochlear sections showed that *Ebf1* deletion increased the number of MYO7A-positive hair cells as well as SOX2-positive supporting and Kölliker’s organ cells from the basal to the apical region at E18.5 (Fig. 3A). An increase in the number of SOX2-positive cells within the medial region of the cochlear floor was also observed at E14.5 (arrowhead in Fig. 1D). In contrast, *Ebf1* deletion had no morphological effects on the vestibular sensory epithelium (Supplemental Fig. S2). Additionally, the apical region of *Ebf1^-/-^* mouse cochleae contained multiple layers of SOX2-positive cells (asterisk in Fig. 3A). IHC analysis of whole-mount cochlear samples showed that *Ebf1^-/-^* mice had an increased number of MYO7A-positive hair cells (Fig. 3B) and ectopic MYO7A-positive cells within the GER (arrows in Fig. 3B). Although the normal cochlea has one and three rows of inner and outer hair cells, respectively, the mutant cochlea had eight to nine rows of hair cells. We found ectopic hair cells in 7 of the 12 examined cochleae. These ectopic MYO7A-positive cells contained stereocilia-like structures, as indicated by phalloidin staining (arrows in Fig. 3C). Increased numbers of supporting cells were confirmed in whole-mount cochlear samples by IHC staining of SOX2 (Fig. 3D), a supporting and Kölliker’s organ cell marker, and PROX1 (Fig. 3E), a pillar and Deiters’ cell marker (Bermingham-McDonogh et al., 2006).

To quantify the number of hair and supporting cells, we counted the cells in three regions within the cochlea (basal, middle, and apical regions; Fig. 4A) and measured the number of MYO7A- or PROX1-positive cells per 200 μm at E18.5. For SOX2-positive cells, we counted the cell number per 100 μm only in the basal regions at E18.5. The results showed that the cochlear hair cell number was significantly increased in *Ebf1^-/-^* mice compared to *Ebf1^+/+^*mice in all three regions (Fig. 4B). *Ebf1^+/-^* mice exhibited a significantly higher number of cochlear hair cells than *Ebf1^+/+^* mice in the middle and apical regions (Fig. 4B, *p* < 0.0001). The hair cell numbers per 200 μm of *Ebf1^+/+^*, *Ebf1^+/-^*, and *Ebf1^-/-^* mice were 142.3 ± 9.0, 150.0 ± 2.8, and 367.5 ± 30.8 in the basal regions; 152.8 ± 8.1, 190.3 ± 7.0, and 363.0 ± 22.4 in the middle regions; and 154.5 ± 5.1, 183.5 ± 6.8, and 440.3 ± 23.8 in the apical regions, respectively.

**Fig. 4.**
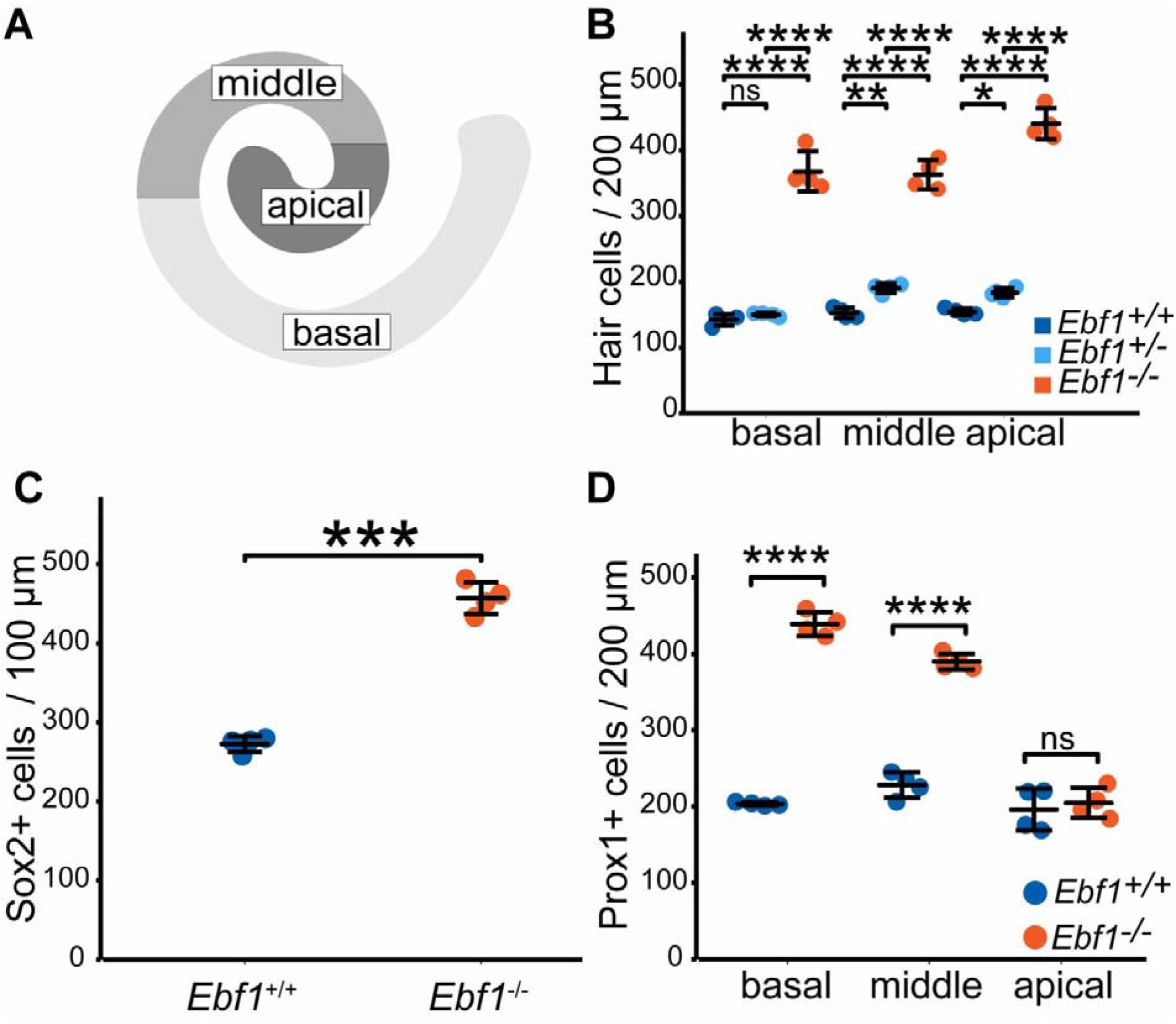
Increases in the number of hair and supporting cells. **A**. Schematic diagram of the cochlea duct showing the positions of the basal, middle, and apical regions of the cochlea. **B**. Total hair cell numbers per 200 μm in the basal, middle, and apical regions of the cochlear ducts of *Ebf1*^+/+^, *Ebf1*^+/-^, and *Ebf1*^-/-^ mice at embryonic day (E) 18.5. **C**. Numbers of SOX2-positive cells per 100 μm in the basal region of the cochlea. **D**. Numbers of PROX1-positive cells per 200 μm in the basal, middle, and apical regions of the cochlea. Two-way analysis of variance (ANOVA) test with Bonferroni *post-hoc* test (**B** and **D**) and Student’s *t*-test (**C**) were performed. \**p* < 0.05, \*\**p* < 0.01, \*\*\**p* < 0.001, and \*\*\*\**p* < 0.0001; ns, not significant. Error bars represent mean ± standard deviation (n = 4).

SOX2-positive cells, constituting a part of Kölliker’s organ cells and supporting cells within organs of Corti, also increased in number by 1.7 times in *Ebf1^-/-^* mice compared to *Ebf1^+/+^* mice (Fig. 4C). PROX1-positive cell numbers significantly increased in *Ebf1^-/-^* mice compared to *Ebf1^+/+^* mice only in the basal and middle regions (Fig. 4D). In the apical region, the number of PROX1-positive cells was similar to that in *Ebf1^-/-^* and *Ebf1^+/+^* mice. The numbers of PROX1-positive cells in *Ebf1^+/+^* and *Ebf1^-/-^* mice were 203.3 ± 2.2 and 439.0 ± 15.4, 228.0 ± 16.8 and 389.8 ± 10.4, and 196.0 ± 27.3 and 204.5 ± 19.6 in the basal, middle, and apical regions, respectively.

We performed IHC analysis using more specific markers to reveal which population of hair or supporting cells increased in number in *Ebf1^-/-^*mice (Supplemental Fig. S3). We immunostained whole-mounted cochlea at E18.5 with anti-VGLUT3 and anti-p75 antibodies, which indicate inner hair (Li et al., 2018) and pillar cells (von Bartheld et al., 1991), respectively. VGLUT3-positive inner hair cells, which are arranged in a single row in wild-type mice, were found to be increased in number in *Ebf1^-/-^* mice (Supplemental Fig. S3A), indicating an increase in both inner and outer hair cells. The number of p75-positive pillar cells, which separate inner and outer hair cells, did not increase in *Ebf1^-/-^* mice (Supplemental Fig. S3B). Considering that PROX1-positive cells indicate pillar and Deiters’ cells, the number of Deiters’ cells increased in *Ebf1^-/-^* mice. However, the arrangement of pillar cells was disrupted in *Ebf1^-/-^* mice (arrows in Supplemental Fig. S3B), which was reflected in the appearance of VGLUT3 cells in the outer hair cell region of *Ebf1^-/-^* mice (arrowheads in Supplemental Fig. S3A).

The total cochlear length, measured based on the length of the MYO7A-positive region (Supplemental Fig. S4A), was slightly, but significantly, shorter in *Ebf1^-/-^* mice than in *Ebf1^+/+^* and *Ebf1^+/-^* mice (Supplemental Fig. S4B, *p* < 0.001).

### Ebf1 deletion caused the aberrant spiral ganglion and nerve fibers

As *Ebf1* is expressed in the spiral ganglion and the number of hair cells, a target of the spiral ganglion cell axon, increased in *Ebf1^-/-^*mice, we examined the spiral ganglion morphology and innervation of cochlear hair cells with IHC using anti-Tubulin β 3 (TUJ1) antibodies at E18.5 (Supplemental Fig. S5).

Compared with *Ebf1^+/+^* mice, which exhibited axons extending from the spiral ganglion cells to the cochlear hair cells, *Ebf1^-/-^* mice had spiral ganglion cells (sg in Supplemental Fig. S5A) under the organs of Corti (arrows in Supplemental Fig. S5A), as well as in their normal position. The axons, which usually run parallel to the rows of outer hair cells, formed a reticulation within the *Ebf1^-/-^* mouse cochlear hair cell regions (Supplemental Fig. S5B). Moreover, the innervation reached Kölliker’s organ (arrowheads and brackets in Supplemental Fig. S5A) as well as the organ of Corti.

### Ebf1 deletion changed the distribution of JAG1-positive Kölliker’s organ cells, the differentiation timing of a prosensory domain, and the proliferation of SOX2-positive cells

As *Ebf1^-/-^*mice had increased numbers of cochlear hair cells, which were differentiated from the prosensory domain, we investigated its specification, differentiation, proliferation, and cell death in the *Ebf1^-/-^* mouse cochlear duct floor.

First, to determine whether formation of the prosensory domain was affected by *Ebf1* deletion, we examined the formation of regions medial and lateral to the prosensory domain. Because these regions express FGF10 and BMP4 to induce non-sensory or sensory epithelia in the cochlear floor (Ohyama et al., 2010; Urness et al., 2015), respectively, we performed ISH for *Fgf10* and *Bmp4* in the basal region of the cochlear duct of *Ebf1^+/+^ and Ebf1^-/-^* mice at E13.5 (Fig. 5A). The formation of both regions was similar in *Ebf1^+/+^* and *Ebf1^-/-^* mice, suggesting that *Ebf1* is not involved in the development of cell populations expressing *Fgf10* or *Bmp4*. To verify the medial cell population more precisely, we immunostained E13.5 and E14.5 cochleae with an anti-JAG1 antibody (Fig. 5A), as JAG1 is exclusively expressed in Kölliker’s organ, a part of the medial region, at this stage (Ohyama et al., 2010). JAG1-positive cells in *Ebf1^-/-^* mouse cochlea expanded to the more medial region compared with those in *Ebf1^+/+^* mouse cochlea at E13.5 and E14.5 (arrowheads in Fig. 5A).

**Fig. 5.**
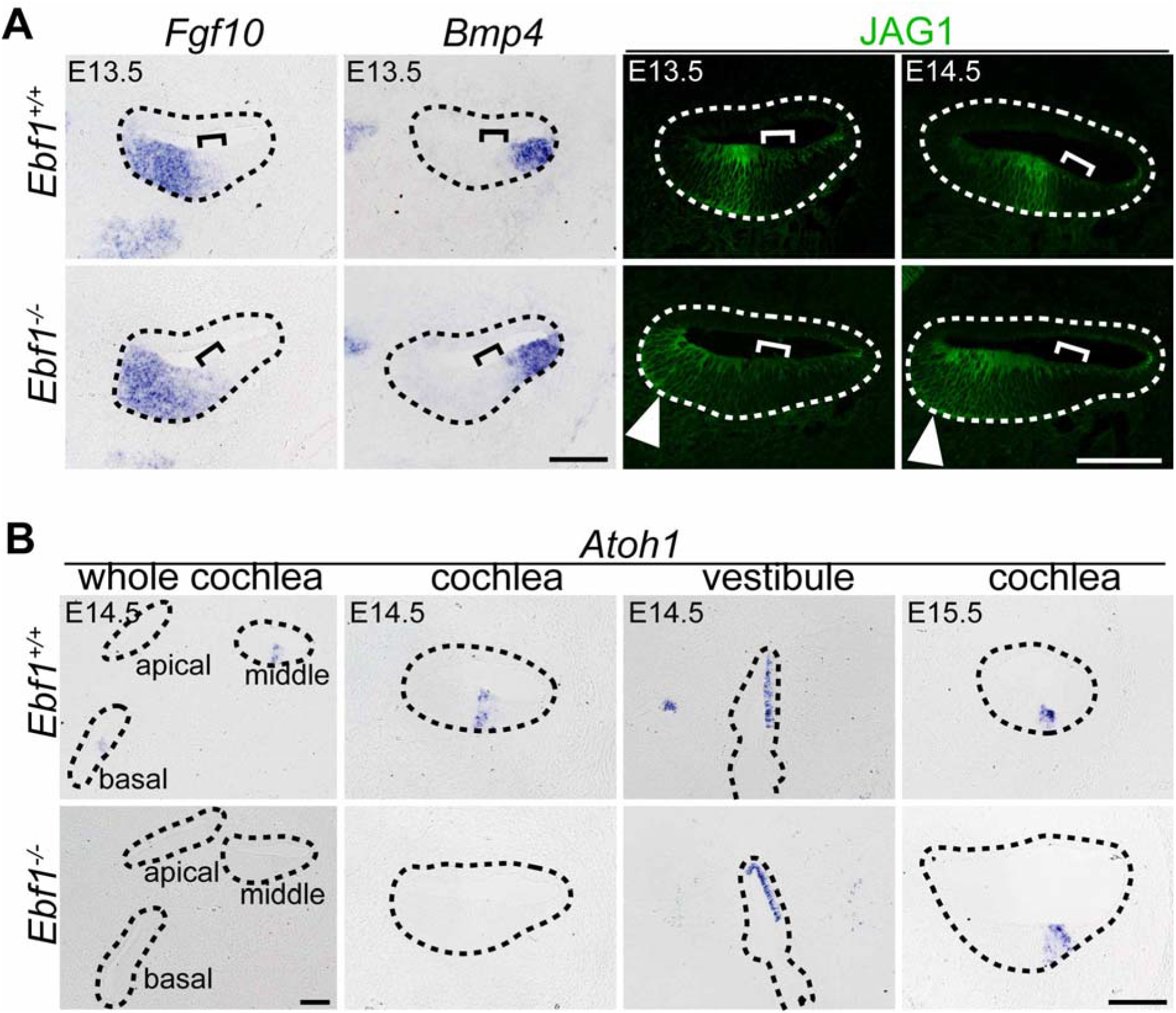
*Ebf1* deletion causes JAG1 expression to spread inward, delaying *Atoh1* expression during cochlear development. **A.** Results of *in situ* hybridization of *Fgf10* and *Bmp4* and immunostaining of JAG1 (green) on cross sections of the basal-to-middle region of the cochlea of *Ebf1^+/+^* and *Ebf1^-/-^* mice at embryonic day (E) 13.5 and E14.5. The prosensory domain is shown in brackets. **B**. Result of *in situ* hybridization of *Atoh1* on cross sections of the basal-to-middle region of the cochlea of *Ebf1^+/+^* and *Ebf1^-/-^* mice at E14.5 and E15.5. Areas enclosed by dashed lines indicate cochlear ducts and vestibules. Scale bars: 100 μm.

Subsequently, we examined the expression of *Atoh1* within the prosensory domain via ISH at E14.5 and E15.5 (Fig. 5B), as *Atoh1* is necessary for hair cells to differentiate from the prosensory cell population (Bermingham et al., 1999) and its expression indicates the initiation of hair cell development from the prosensory domain. Compared to *Ebf1^+/+^* mice that expressed *Atoh1* within the prosensory domain from E14.5, *Atoh1* mRNA was not detected in the basal to middle region of the E14.5 *Ebf1^-/-^* mouse cochlea, although the vestibular organs expressed *Atoh1* within the prosensory epithelia. However, E15.5 *Ebf1^-/-^*mice exhibited an *Atoh1* signal within the prosensory domain of the cochlea. This result suggests that while the cell fate specification of sensory epithelia in the cochlea is not affected, its timing is delayed by *Ebf1* deletion. Considering that the differentiation of cochlear sensory epithelia promotes the transition from the basal to apical turns of the cochlea (Sher, 1971), the expression of hair and supporting cell markers at a later stage, E18.5, also indicated delayed differentiation of *Ebf1^-/-^*mouse cochleae (Supplemental Fig. S3). These markers were found to be detected in more basal cochlear regions in *Ebf1^-/-^* mice than in *Ebf1^+/+^* mice.

The expansion of the JAG1-positive cell area and the increased numbers of hair and supporting cells suggest that the enhancement of proliferation or suppression of cell death occurs within the prosensory domain and Kölliker’s organ of *Ebf1^-/-^* mouse cochlea. To identify the mechanisms that correlate with the functions of EBF1 within the cochlea, we tested the proliferation and apoptotic status of *Ebf1^-/-^* mouse cochlea. To evaluate the proliferation status of the prosensory domain and Kölliker’s organ, we immunostained cochlear sections with SOX2, a marker of the prosensory domain and a part of Kölliker’s organ, and 5-ethynyl-2′-deoxyuridine (EdU) at E12.5, 13.5, 14.5, and 16.5 after administering EdU to pregnant mice (Fig. 6A). Quantification of SOX2-positive cells showed that their number decreased in *Ebf1^+/+^* mice from E12.5 onward (Fig. 6B), and their location was limited to the prosensory domain (brackets in Fig. 6A). In contrast, SOX2-positive cells in *Ebf1^-/-^* mouse cochlea were found both in the prosensory domain and the medial region, even at E13.5, which was consistent with the results of JAG1 immunostaining (Fig. 5A). The number of SOX2-positive cells in *Ebf1^-/-^* mice was similar to that in *Ebf1^+/+^* mice at E12.5, but increased at E13.5 and returned to the E12.5 level at E14.5. Therefore, the number of SOX2-positive cells in *Ebf1^-/-^* mice was significantly higher than those in *Ebf1^+/+^*mice at E13.5 and E14.5 (Fig. 6B). The number of EdU-positive proliferating cells within SOX2-positive cells was significantly higher in *Ebf1^-/-^* mice than in *Ebf1^+/+^* mice at E13.5 and E14.5 (Fig. 6C). As the number of SOX2-positive cells increased in *Ebf1^-/-^* mice after E13.5 (Fig. 6B), normalization of SOX2-positive cell numbers was necessary to correctly evaluate the proliferation status of SOX2-positive cells. We calculated the proportion of EdU-positive cells among SOX2-positive cells and found that the proliferation was enhanced in the SOX2-positive cells of *Ebf1^-/-^*mouse cochlea only at E13.5 (49.8 ± 4.3%) compared with that in *Ebf1^+/+^*mouse cochlea (37.8 ± 4.2%) (Fig. 6D). These results suggested that EBF1 suppressed the proliferation of SOX2-positive cells within a limited time window. Morphologically, a difference in proliferation was observed in the prosensory domain, as indicated by EdU immunostaining (brackets at E13.5; Fig. 6A). EdU staining was observed in the Kölliker’s organs of *Ebf1^-/-^*mice, even at E16.5, but not observed in *Ebf1^+/+^* mice (E16.5 of Fig. 6A).

**Fig. 6.**
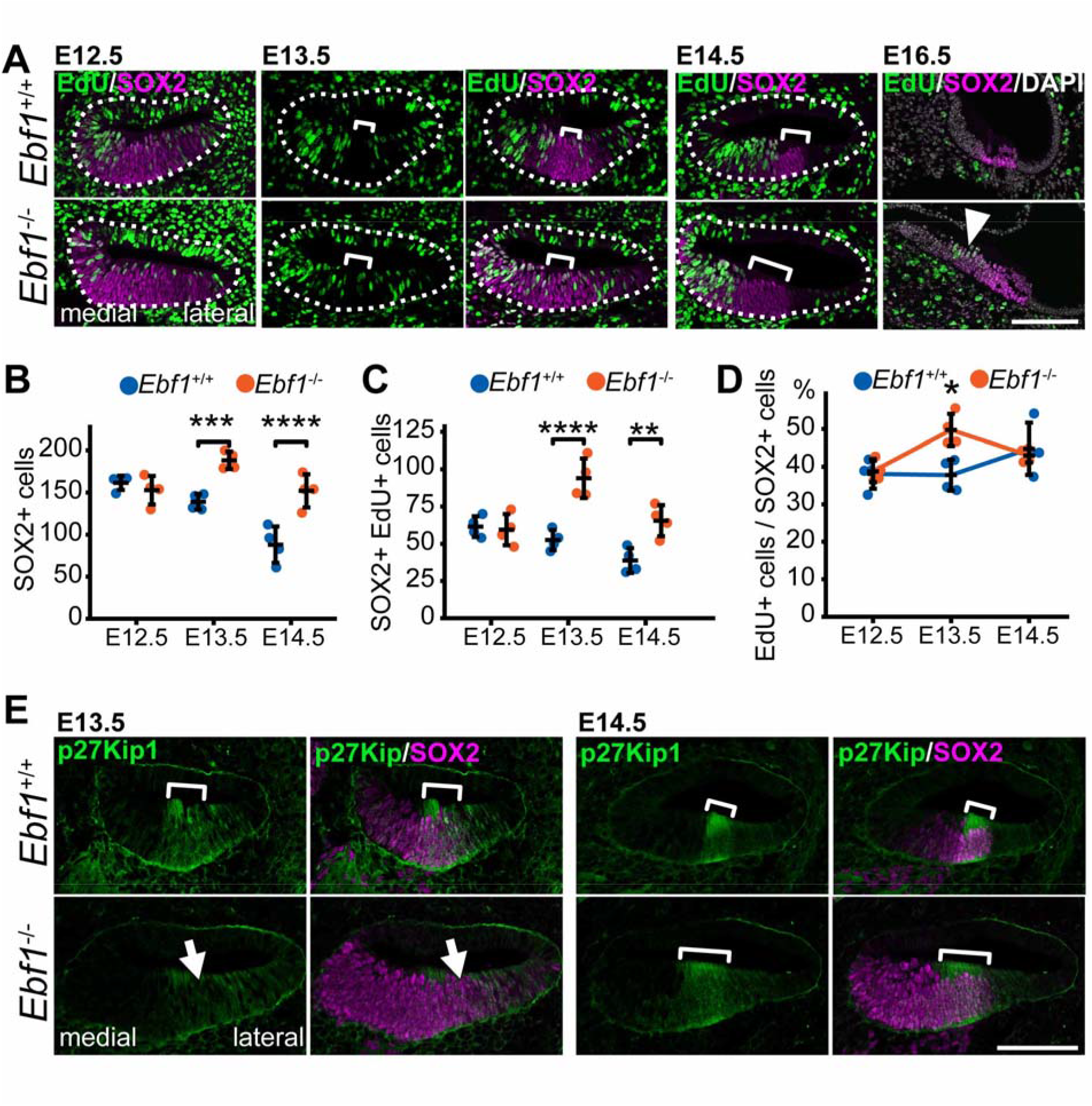
Effect of *Ebf1* deletion on cell proliferation during inner ear development. **A**. Cross sections of the cochlear basal regions at embryonic day (E) 12.5, E13.5, E14.5, and E16.5 from *Ebf1^+/+^* and *Ebf1^-/-^*mice. Sections were immunostained with 5-ethynyl-2’-deoxyuridine (EdU, green) and SOX2 (magenta). E16.5 sections were counter-stained with 4’,6-diamidino-2-phenylindole (DAPI, gray). Areas enclosed by dashed lines indicate the cochlear ducts and brackets indicate prosensory domains. **B**–**D**. Quantitative assessment of the SOX2-positive region in the cochlea epithelia. The numbers of SOX2-positive cells (**B**) as well as EdU- and SOX2-double positive cells (**C**) were counted, and the percentage of EdU-positive cells among SOX2-positive cells (**D**) was calculated. \**p* < 0.05, \*\**p* < 0.01, \*\*\**p* < 0.001, and \*\*\*\**p* < 0.0001. Error bars represent mean ± standard deviation. n = 4. **E**. Cross sections of the cochlear basal regions at E13.5 and E14.5 from *Ebf1^+/+^* and *Ebf1^-/-^*mice immunostained with p27Kip1 (green) and SOX2 (magenta). Scale bars: 100 μm.

The loss of proliferation within the prosensory domain around E13.5 (bracket in the *Ebf1^+/+^* sample at E13.5, Fig. 6A) has been well documented in previous studies (Chen and Segil, 1999; Chen et al., 2002). The post-mitotic domain is called the zone of non-proliferating cells (ZNPC), and is characterized by the expression of the cyclin-dependent kinase inhibitor p27Kip1. To confirm the EdU immunostaining results, we performed p27Kip1 immunostaining at E13.5 and E14.5 (Fig. 6E). Although p27Kip1 was detected in the prosensory domain of *Ebf1^+/+^* mice at E13.5, it was not expressed in *Ebf1^-/-^* mouse cochlea at this stage (arrows in Fig. 6E), which was consistent with the results of EdU detection. At E14.5, p27Kip1 was detected in a larger area of the middle part of *Ebf1^-/-^* mouse cochlear floors than in those of *Ebf1^+/+^*mice.

In contrast, apoptosis within the inner ear or cochlear duct did not increase in *Ebf1^-/-^* mice at E11.5 and E13.5, compared with that in *Ebf1^+/+^*mice (Supplemental Fig. S6).

### Ebf1 deletion impairs auditory function

The aberrant cochlear sensory epithelia observed in *Ebf1*-deleted mice suggest that hearing ability is impaired in these mice. To evaluate the effect of *Ebf1* deletion on auditory function, we measured the auditory brainstem response (ABR) (Fig. 7A) and distortion product of otoacoustic emissions (DPOAE) (Fig. 7B) in P21 *Foxg1Cre;Ebf1^fl/fl^* mice. We did not use *Ebf1^-/-^* mice for this analysis to avoid embryonic lethality and to eliminate the effects of the hypoplastic scala tympani observed in *Ebf1^-/-^*mice on auditory function. The phenotype of *Foxg1Cre;Ebf1^fl/fl^* mice was evaluated using whole-mount cochlear samples collected at P23 (Fig. 7C). We observed a marked increase in the number of cochlear hair cells in *Foxg1Cre;Ebf1^fl/fl^* mice, comparable to the morphology of E18.5 *Ebf1^-/-^* mice (Fig. 3B and Fig. 7C).

**Fig. 7.**
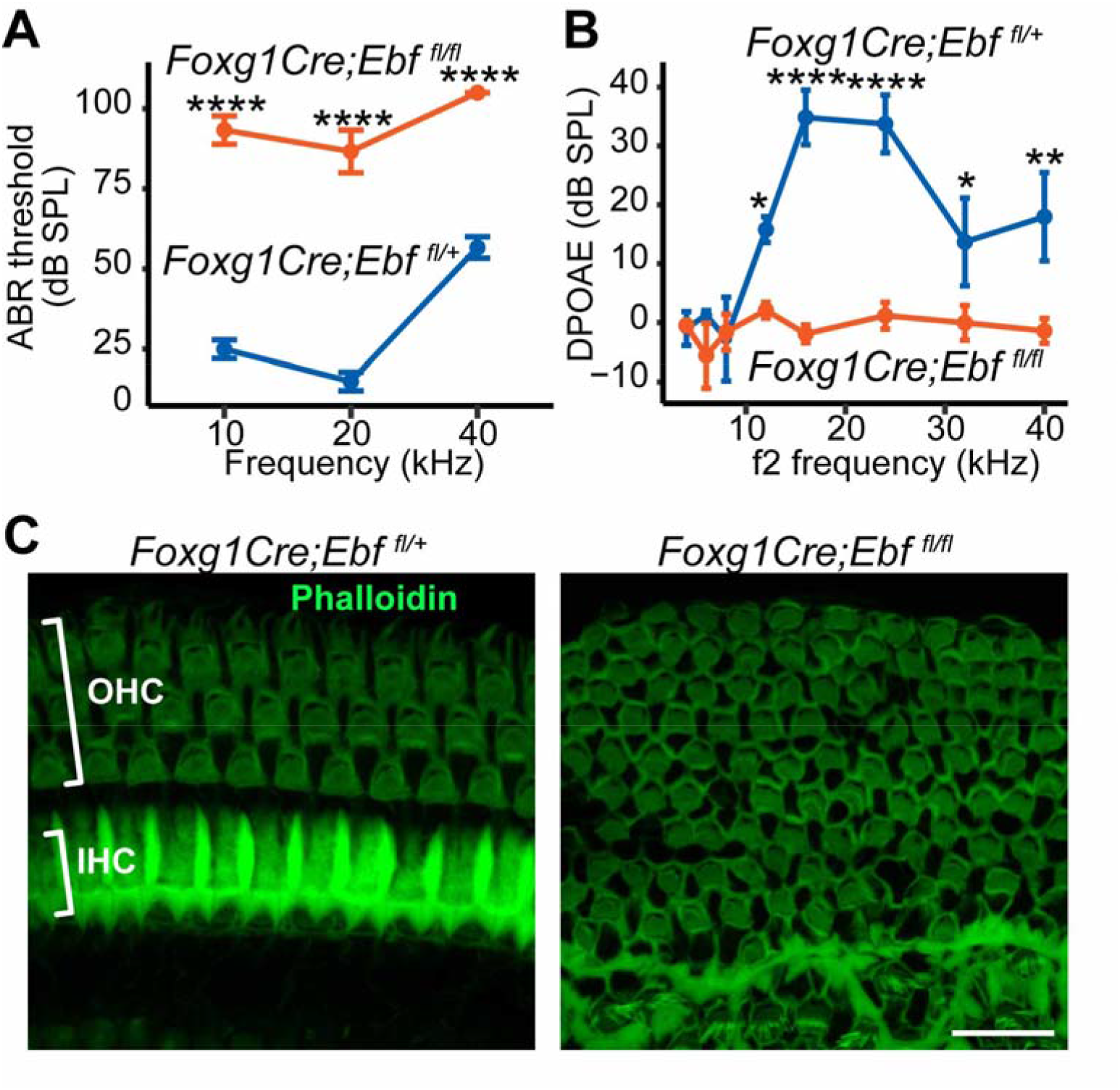
*Ebf1* deletion impairs auditory function. **A**. Auditory brainstem response (ABR) thresholds of P21 *Foxg1Cre;Ebf1^fl/fl^*(red) and *Foxg1Cre;Ebf1^fl/+^* (blue, control) mice. **B**. Distortion product optoacoustic emissions (DPOAE) were measured at P21 from *Foxg1Cre;Ebf1^fl/fl^*(red) and control (blue) mice. \**p* < 0.05, \*\**p* < 0.01, and \*\*\*\**p* < 0.0001 from one-way analysis of variance (ANOVA) with Tukey’s HSD *post-hoc* test. Error bars represent mean ± standard error of mean. **C**. The whole mount images of the cochlear basal regions from *Foxg1Cre;Ebf1^fl/+^* and *Foxg1Cre;Ebf1^fl/fl^* mice at P23 labeled with phalloidin (green). Scale bar: 20 μm.

ABR measurement showed significant elevations of thresholds of the response to sound in *Foxg1Cre;Ebf1^fl/fl^* mice at all frequencies examined (10 kHz, 93.3 ± 2.5 dB; 20 kHz, 86.7 ± 3.8 dB; 40 kHz, 105.0 ± 0.0 dB) (Fig. 7A) compared with control mice, indicating severe hearing loss in *Ebf1*-deleted mice. Subsequently, we performed DPOAE tests to assess the function of the increased number of outer hair cells caused by *Ebf1* deletion because DPOAE detects nonlinear responses of outer hair cells to sound. The DPOAE responses in *Foxg1Cre;Ebf1^fl/fl^*mice were significantly lower than those in control mice (Fig. 7B). The decreased DPOAE response in *Foxg1Cre;Ebf1^fl/fl^* mice also suggests that the increased number of hair cells caused by *Ebf1* deletion did not function as outer hair cells.

## Discussion

The results of this study indicate a novel and interesting role of *Ebf1* in the development of the cochlear epithelia and otic mesenchyme. *Ebf1* is involved in the formation of appropriate numbers of hair and supporting cells within the cochlear sensory epithelia. Moreover, *Ebf1* is important for the development of the scala tympani, which is formed through *Ebf1* expression within the mesenchyme, and spiral ganglion cells. The *Ebf1* deficiency in the cochlear epithelia resulted in impaired auditory function, suggesting that *Ebf1* is necessary for normal auditory function.

Since EBF1 was originally identified from the regulators of early B cell differentiation (Hagman et al., 1991) and olfactory-specific genes (Wang and Reed, 1993), the expression patterns and functions of EBF1 have been studied in B lymphocytes (Lin and Grosschedl, 1995), olfactory epithelia (Wang et al., 1997), striatum neurons (Garel et al., 1999), the retina (Jin et al., 2010; Jin and Xiang, 2011), the kidney (Fretz et al., 2014; Nelson et al., 2019), bone marrow stromal cells (Derecka et al., 2020), osteoblasts (Hesslein et al., 2009), and adipose tissue (Jimenez et al., 2007). The expression of *Ebf1* and its protein product has been reported in all three germinal layers, and the function of EBF1 varies (Liberg et al., 2002). EBF1 is involved in cell fate specification (Jin et al., 2010; Nechanitzky et al., 2013), cell differentiation (Lin and Grosschedl, 1995; Wang et al., 1997; Garel et al., 1999; Jimenez et al., 2007; Jin and Xiang, 2011; Nieminen-Pihala et al., 2021), cell maturation (Fretz et al., 2014; Nelson et al., 2019), cell migration (Derecka et al., 2020), and path finding by neurons (Garel et al., 2000; Jin and Xiang, 2011), mainly during the developmental stage in mammals.

The present study revealed that *Ebf1* is expressed not only in the inner ear epithelium and spiral ganglion but also in the otic mesenchyme. As the inner ear epithelium and spiral ganglion cells are derived from the ectoderm (Wu and Kelley, 2012), *Ebf1* is expressed in both ectodermal and mesenchymal tissues within the inner ear. Its expression in the inner ear began at approximately E10.5, as confirmed by qRT-PCR and ISH (Fig. 1A and C). Within the inner ear epithelia, *Ebf1* showed different expression patterns in the cochlear and vestibular organs from E13.5 to E18.5. In the vestibule and crista, *Ebf1* was expressed in parts of *Sox2*-positive prosensory regions (Fig. 1B, black arrows and C). However, its expression in the cochlea was not limited to a part of the *Sox2*-positive prosensory domain (white arrows in Fig. 1B) but expanded toward a more medial region in the cochlear floor, where Kölliker’s organ exists (Fig. 1B–D). Thus, the *Ebf1* expression area comprised most of the GER. This difference suggests distinct functions of *Ebf1* in the development of cochlear and vestibular organs.

To elucidate the function of *Ebf1* in inner ear development, we examined the inner ear morphology of *Ebf1* conventional and inner ear-specific conditional knockout mice. As suggested by the expression study, the cochlear and vestibular organs exhibited different morphological phenotypes. No differences were observed in the vestibular morphology of *Ebf1^+/+^*and *Ebf1^-/-^* mice (Supplemental Fig. S2), despite *Ebf1* expression. As observed in the olfactory epithelia and retina (Wang and Reed, 1993; Wang et al., 1997; Jin et al., 2010), the deletion of *Ebf1* may be compensated for by other *Ebf* family genes, and studies in other mutant mice, including double- or triple-knockout mice and mice with the dominant negative form of the EBF protein, are needed to elucidate the function of EBF1 in the vestibular organ.

In contrast, the cochlea of *Ebf1*-deleted mice showed various phenotypes, indicating that other *Ebf* subtypes do not have redundant functions with *Ebf1* in the cochlea as in B cells, osteoblasts, and the striatum (Lin and Grosschedl, 1995; Garel et al., 1999; Nieminen-Pihala et al., 2021). HE staining revealed loss of the scala tympani and spiral limbus (Fig. 2C). Because both structures are derived from mesenchymal tissues (Sher, 1971; Phippard et al., 1999), we hypothesized that these phenotypes reflect the roles of EBF1 in cochlear mesenchymal tissue. To confirm this, we compared the formation of the scala tympani and spiral limbus in two *Ebf1*-deleted mouse strains, conventional (*Ebf1^-/-^*mice) and epithelia-specific conditional (*Foxg1Cre;Ebf1^fl/fl^* mice) *Ebf1* knockout mice (Fig. 2C). Although the scala tympani developed normally in *Foxg1Cre;Ebf1^fl/fl^*mice, the spiral limbus was hypoplastic in both *Foxg1Cre;Ebf1^fl/fl^*and *Ebf1^-/-^* mice. These results clearly indicate that EBF1 in the cochlear mesenchyme is involved in the formation of the scala tympani. In contrast, spiral limbus formation depends on both mesenchymal and epithelial or epithelial expression of *Ebf1*. Because *Ebf1* was expressed in both the epithelia and mesenchyme (Fig. 1B and Supplemental Fig. S1), we were unable to determine which was the case in this study; however, mesenchyme-specific deletion of *Ebf1* will confirm the tissue responsible for inducing the spiral limbus. Gross morphological observation (Fig 2B) and MYO7A staining of the whole-mount cochlea (Supplemental Fig. S4) showed that the length of the cochlear duct was shorter in *Ebf1^-/-^* mice. As the number of cochlear turns in the cochlear section was lower in *Ebf1^-/-^* mice than in *Foxg1Cre;Ebf1^fl/fl^*mice, this phenotype reflects the function of mesenchymal EBF1 in regulating the length of the cochlear duct.

In addition to the mesenchyme, more prominent roles of *Ebf1* have been found in the cochlear epithelia. In both *Ebf1^-/-^* and *Foxg1Cre;Ebf1^fl/fl^*mice, the numbers of both hair and supporting cells increased throughout cochlear turns at E18.5 (Fig. 3 and 4). We observed an increase in the numbers of both inner and outer hair cells (Supplemental Fig. S3A). However, more detailed IHC analysis of supporting cells showed that only the Deiters’ cells increased in number among PROX1-positive and SOX2-positive cells located medial to the organs of Corti, and the number of pillar cells did not increase (Supplemental Fig. S3B). This phenotype suggests that EBF1 is involved in the regulation of hair and supporting cell number during cochlear development. To determine the mechanisms of hair and supporting cell number increase under *Ebf1* knockout conditions, we evaluated the specifications and differentiation of the cochlear prosensory domain and its proliferation and cell death status. We found that the formation of cochlear non-sensory regions medial and lateral to the prosensory domain were normal in *Ebf1^-/-^*mice (Fig. 5A). Although the lateral region marker BMP4 has been reported to affect the development of the prosensory domain (Ohyama et al., 2010), the normal formation of this region indicates that the phenotypes of *Ebf1^-/-^* mouse cochlear sensory epithelia were not caused by aberrant BMP4 expression, but by factors within the prosensory domain. In contrast to markers outside the prosensory domain, the molecules expressed in the prosensory domain and Kölliker’s organs, JAG1 and SOX2, showed abnormal expression patterns (Fig. 1D, Fig. 3A, and Fig. 5A). These two molecules were expressed in a more medial region of the *Ebf1^-/-^*mouse cochlear floor at E14.5 and E18.5. The fact that EBF1 was expressed in a more medial region than SOX2 in wild-type mice indicates that it plays a role in suppressing the localization of JAG1- and SOX2-positive cells in the most medial region. The study of proliferation status within the SOX2-positive cells showed that *Ebf1* deletion enhanced the proliferation of SOX2-positive cells specifically at E13.5 (Fig. 6 D), which was supported by the loss of p27Kip1 expression in the possible prosensory domain of the *Ebf1^-/-^*mice at E13.5 (Fig. 6E). The highest *Ebf1* expression level at E13.5 (Fig. 1A) may be related to these phenotypes in *Ebf1^-/-^* mice. This aberrant proliferation within SOX2-positive cells was suggested to increase the numbers of hair and supporting cells at later stages (Fig. 3A, B, D, and E). Evaluation of hearing ability at the postnatal stage showed that an increase in hair and supporting cell numbers resulted in an increased hearing threshold (Fig. 7). These results indicate that EBF1 suppresses the proliferation of SOX2-positive cells and thus contributes to the development of appropriate numbers of hair and supporting cells, resulting in the development of normal auditory function. Because the SOX2-positive cells within the E13–14 cochlea become Kölliker’s organ as well as lateral and medial prosensory domains at a later stage (Kolla et al., 2020) and the GER has the potential to form cochlear sensory epithelia (Kubota et al., 2021), EBF1 expressed in SOX2-positive cells regulates the number of hair and supporting cells. Several lines of evidence support the function of EBF1 to suppress cell proliferation. Human EBF1 has been reported to suppress the proliferation of malignant tumors (Shen et al., 2020), and the deletion of *Rb1*, a known tumor suppressor and cell cycle regulator (Lipinski and Jacks, 1999; Classon and Harlow, 2002), results in the same morphology in the cochlea as that caused by *Ebf1* deletion (Sage et al., 2005).

The expression of mature cochlear cell markers, including MYO7A, VGULT3, and p75, in *Ebf1^-/-^* mice indicated that each cell type developed with normal cell fate specification. Although some GER cells in *Ebf1^-/-^* mice expressed the hair cell marker MYO7A (Fig. 3B and C) and cell fate specification appeared to deteriorate, the penetrance of this phenotype was low. Cell fate may be regulated by EBF1 in the cochlea to a small extent; however, cell specification does not play a prominent role in cochlear EBF1, which is different from other tissues, including B lymphocytes (Nechanitzky et al., 2013). In contrast, several results from our study indicate that the speed of differentiation appears to deteriorate in the *Ebf1^-/-^* mouse cochlea. The expression of *Atoh1* within the cochlear prosensory domain, which was observed at E13.5 in wild-type mice, was detected as late as E14.5 in *Ebf1^-/-^* mice (Fig. 5B). The regions where VGLUT3- and p75-positive cells were detected in *Ebf1^-/-^* mouse cochlea were more basal than those in *Ebf1^+/+^*mouse cochlea at E18.5 (Supplemental Fig. S3). This delay in differentiation may be caused by the aberrant proliferation of the prosensory domain in *Ebf1^-/-^* mice, as the deterioration of proliferation affects the differentiation of cells (Ruijtenberg and van den Heuvel, 2016).

In addition to the cochlear epithelia and mesenchyme, deletion of *Ebf1* resulted in an altered neuroaxonal composition of spiral ganglion neuronal cells in the organ of Corti, suggesting that EBF1 may also affect the pathfinding of spiral ganglion cells within the cochlea, as observed in the aberrant pathfinding of facial branchiomotor neurons and retina (Garel et al., 2000; Jin and Xiang, 2011). The present study was limited to IHC analysis of TUJ1, and further studies are needed to elucidate the role of *Ebf1* in spiral ganglion differentiation and development. Spiral ganglion axons synapse with hair cells from within the otic mesenchyme, and axons extend through the mesenchyme to innervate the organ of Corti (Coate and Kelley, 2013). Thus, deletion of *Ebf1* in the mesenchyme may also contribute to the phenotype observed in the spiral ganglion cells.

The expression of *Ebf1* in the medial region of the cochlear floor (Fig. 1B, C, and D) and the spiral limbus loss (Fig. 2C) and ectopic MYO7A-positive cells within the GER (Fig. 3B and C) in *Ebf1*-deleted mice suggest that EBF1 has functions within the GER or Kölliker’s organ. Recently, PRDM16 was reported to be a marker of Kölliker’s organ and an important molecule in its development (Ebeid et al., 2022). *Prdm16* null mutant mice showed phenotypes similar to those of *Ebf1*-deleted mice, including shortened cochlear length, spiral limbus loss, and ectopic hair cells within the GER. Moreover, the *Prdm16* null mutant mouse cochlear duct expressed approximately half the amount of *Ebf1* compared to the wild-type mouse cochlear duct (Ebeid et al., 2022), suggesting that EBF1 is involved in the development of Kölliker’s organ. However, if PRDM16 is the direct upstream regulator of EBF1, the *Ebf1* expression level should be much less in *Prdm16* null mutant mice, and these mice should exhibit increased hair and supporting cells and loss of scala tympani, as observed in *Ebf1^-/-^* mice. Thus, PRDM16 does not control the expression of EBF1.

The phenotypes of the increased numbers of hair and supporting cells suggest the involvement of previously reported molecules crucial for the development of cochlear sensory epithelia, including Notch signal-related molecules (Yamamoto et al., 2011) and SOX2 (Kiernan et al., 2005), in the regulation of *Ebf1* expression. However, *Foxg1Cre*-mediated *Notch 1*- or *Sox2*-deleted mice do not show lower *Ebf1* expression levels in striatal neurons (Mason et al., 2005; Ferri et al., 2013), suggesting that Notch signaling and SOX2 do not control EBF1 expression.

Identification of the molecules upstream and downstream of EBF1 will be the next step in revealing the precise function of EBF1 in the cochlea and the grand scheme of inner ear development.

In conclusion, our study confirms that *Ebf1* and its protein are expressed in the epithelia of the inner ear prosensory domain as well as in Kölliker’s organ, the mesenchyme, and CVG cells within the cochlea, and play important roles in the formation of each structure. Epithelial EBF1 regulates the number of cochlear hair and supporting cells by suppressing the proliferation of the prosensory domain and Kölliker’s organ cells, mainly at E13.5, whereas mesenchymal EBF1 contributes to scala tympani formation. Therefore, epithelial EBF1 is crucial for normal hearing in mammals.

## Material and Methods

### Animals

Slc: ICR mice were purchased from Japan SLC (Hamamatsu, Japan). *Ebf1^-/-^*mice (Lin and Grosschedl, 1995) and *Ebf1^fl/fl^* (Gyory et al., 2012) were used in this study. *Ebf1^fl/+^* mice were crossed with *Foxg1^Cre/+^ mice* (*Foxg1Cre*) (Hébert and McConnell, 2000) and *Ebf1^-/-^* or *Foxg1Cre;Ebf1^fl/fl^*mice were used as experimental animals. Additionally, *Ebf1^+/+^*, *Ebf1^+/-^*, or *Foxg1Cre;Ebf1^fl/+^* mice were used as controls.

*Ebf1^+/-^*, *Foxg1Cre*, and *Ebf1^fl/fl^*mice were maintained on a C57BL/6 background. All experimental protocols were approved by the Animal Research Committee of Kyoto University (Med Kyo 20132, Kyoto, Japan). All animal experiments were performed according to the National Institutes of Health Guidelines for the Care and Use of Laboratory Animals. All the animals used in this study were maintained at the Institute of Laboratory Animals, Graduate School of Medicine, Kyoto University. The mice were mated in the evening, and vaginal plugs were checked early in the morning. The day a vaginal plug was detected was defined as E0.5.

### qRT-PCR

Inner ears were dissected from E9.5, E10.5, E11.5, E12.5, E13.5, E14.5, E16.5, E18.5, and P0 ICR mice. After the surrounding tissue was removed from the inner ears, at least four samples were immersed in TRIzol Reagent (Thermo Fischer Scientific, Waltham, MA, USA) and preserved at −80 ℃ until RNA extraction. Total RNA was extracted using the RNeasy™ Mini Kit (QIAGEN, Venlo, Netherlands) and reverse transcribed using the ReverTra Ace™ qPCR RT Master Mix with gDNA Remover (TOYOBO, Osaka, Japan). The cDNA was mixed with PowerUp SYBR Green Master Mix (Applied Biosystems) and various sets of gene-specific forward and reverse primers and subsequently subjected to real-time PCR quantification using a StepOnePlus Real-Time PCR System (Applied Biosystems, Waltham, MA, USA). The following primer sequences were used: *Ebf1* forward, AACTCCAAGCACGGGCGGAG; *Ebf1* reverse, CGGGCTGATGGCTTTGATACAGG; *Rplp0* forward, CACTGGTCTAGGACCCGAGAAG; *Rplp0* reverse, GGTGCCTCTGGAGATTTTCG. Relative mRNA expression levels were calculated using the standard curve method, and the mouse housekeeping gene *Rplp0* was used as an invariant control.

### ISH

Whole embryos (E9.5–E11.5) and whole heads (E12.5–P0) were fixed in 4% paraformaldehyde (PFA; Nacalai Tesque, Kyoto, Japan) in 0.1 M phosphate-buffered saline (PBS) at 4 ℃ overnight. Samples were cryoprotected in 30% sucrose/PBS (Nacalai Tesque), embedded in Tissue-Tek® O.C.T.™ compound (Sakura Finetek Japan, Tokyo, Japan), and sectioned at 10-μm thickness using a cryostat (CryoStar™ NX70; Thermo Fisher Scientific). The sections were subsequently mounted on silane-coated glass slides (Matsunami Glass, Osaka, Japan).

cDNA fragments were generated by PCR using E13.5 inner ear cDNA of Slc:ICR mice and subsequently cloned into the pCR®-Blunt Ⅱ-TOPO® vector (Invitrogen, Waltham, MA, USA) to prepare RNA probe templates. We synthesized digoxigenin (DIG)-labeled sense and antisense RNA probes using a DIG RNA Labeling Kit (Roche, Basel, Switzerland) after digestion with the appropriate restriction enzymes *Bam*HⅠ-HF, *Hind*Ⅲ, *Not*Ⅰ-HF, *Sac*Ⅰ, or *Xho*Ⅰ (New England Biolabs, Ipswich, MA, USA). The following probes were used for ISH: *Ebf1* (NM_001,290,709, nucleotides 1436–2269), *Sox2* (IMAGE clone: 6413283), *Bmp4* (NM_007554.3, nucleotides 1013-1876), *Atoh1* (NM_007500.5, nucleotides 13-2111), and *Fgf10* (NM_008002.5, nucleotides 571-1027). Each corresponding sense probe was used as a negative control.

Sections were fixed with 4% PFA and 0.2% glutaraldehyde (Nacalai Tesque) in PBS at room temperature (RT) for 10 min, bleached with 6% hydrogen peroxidase (FUJIFILM Wako Pure Chemical Corporation, Osaka, Japan) in 0.1% Tween-20 in PBS (PBST) at RT for 10 min, treated with proteinase K (20 μg/μL, Roche) for 5 min, and re-fixed with 4% PFA and 0.2% glutaraldehyde in PBS at RT for 10 min.

The prehybridization was performed in hybridization solution containing 50% formamide (Nacalai Tesque), 5× saline sodium citrate buffer (SSC, Nacalai Tesque; adjusted to pH 4.5 with citrate), 1% sodium dodecyl sulfate (Sigma-Aldrich, St. Louis, MO, USA), 50 μg/mL yeast RNA (Invitrogen), and 50 μg/mL heparin (Sigma-Aldrich) at 70 ℃ for 1 h. For hybridization, we incubated the sections in a hybridization solution with a 0.2 μg/mL DIG-labelled RNA probe at 70 ℃ for 16 h in sealed plastic bags.

Sections were rinsed first in 50% formamide with 6× SSC and 1% sodium dodecyl sulfate at 70 ℃, then in 50% formamide with 2.4× SSC at 65 ℃, and finally in 1× Tris-buffered saline (Nacalai Tesque) with 0.1% Tween 20 (TBST) at RT. Sections were blocked with 5% sheep serum (Sigma-Aldrich) and incubated with a 1:4000 dilution of Anti-Digoxigenin-AP Fab fragments (Roche) at 4 ℃ overnight.

After rinsing with TBST and NTMT containing 100 mM NaCl (Nacalai Tesque), 100 mM Tris-HCl pH 9.5, 10 mM MgCl_2_ (FUJIFILM Wako Pure Chemical Corporation), 0.1% Tween-20, and 480 μg/mL levamisole (Sigma-Aldrich), the sections were incubated with nitro-blue tetrazolium chloride (Roche) and 5-bromo-4-chloro-3-indolyl phosphate solution (Roche). Images were captured using a BX-50 microscope (Olympus Corp., Tokyo, Japan).

### IHC analysis

IHC sections were prepared in a manner similar to that used for ISH.

After washing with PBS, all samples were incubated with Blocking One Histo (Nacalai Tesque) for 10 min at RT and 10% normal donkey serum (Sigma-Aldrich) in PBS/0.5% Triton X-100 with 5% Blocking One Histo for 30 min at RT.

The samples were stained with primary antibodies at 4 ℃ overnight or RT for 1 h. After washing with PBST, the samples were incubated with Alexa Fluor secondary antibodies. F-actin (actin filaments) was stained with phalloidin 647 (1:500; Thermo Fisher Scientific) at RT for 1 h. Nuclei were stained with 4′,6-diamidino-2-phenylindole (DAPI; Thermo Fisher Scientific).

The following primary antibodies were used in this study: rabbit anti-EBF1 antibody (1:1000, AB10523; Millipore, Darmstadt, Germany,), mouse anti-MYO7A antibody (1:1000, 138-1; Developmental Studies Hybridoma Bank, Iowa City, IA, USA), rabbit anti-MYO7A antibody (1:1000, 25-6790; Proteus BioSciences, Waltham, MA, USA), goat anti-SOX2 antibody (1:250, AF2018; R&D Systems, Minneapolis, MN, USA), rabbit anti-SOX2 antibody (1:100, 11064-1-AP; Proteintech, Manchester, UK), rabbit anti-VGLUT3 antibody (1:500, 135 203; Synaptic systems, Goettingen, Germany), goat anti-JAG1 antibody (1:500, sc-6011; Santa Cruz, Dallas, TX, USA), mouse anti-p27Kip1 antibody (1:200, 610242; BD Transduction Laboratories), rabbit anti-Tubulin β 3 (TUJ1) antibody (1:1000, PRB-435P; Biolegend, San Diego, CA, USA), rabbit anti-PROX1 antibody (1:500, AB5475; Millipore), rabbit anti-p75 antibody (1:500, AB1554; Millipore), and rabbit anti-cleaved caspase 3 (CC3) antibody (1:400, Asp175; Cell Signaling Technology, Danvers, MA, USA).

Antigen retrieval was performed for p27Kip1 staining by heating sections in HistoVT one (Nacalai Tesque) at 90 ℃ for 10 min prior to the addition of the primary antibodies.

The secondary antibodies used were Alexa Flour 488 donkey anti-rabbit IgG, Alexa Flour 488 donkey anti-goat IgG, Alexa Flour 488 donkey anti-mouse IgG, Alexa Flour 568 donkey anti-rabbit IgG, Alexa Flour 568 donkey anti-goat IgG, Alexa Flour 633 donkey anti-rabbit IgG, Alexa Flour 633 donkey anti-goat IgG, and Alexa Flour 633 donkey anti-mouse IgG (1:500; Thermo Fisher Scientific).

The sections were mounted using Fluoromount-G® Anti-Fade (Southern Biotechnology Associates Inc., Birmingham, AL, USA). Images of the specimens were captured using an Olympus BX50 microscope (Olympus), an Olympus DP70 digital camera (Olympus), and a Zeiss LSM900 with Airyscan2 (Carl Zeiss AG, Oberkochen, Germany).

### HE staining

Frozen sections (10-µm thick) were immersed in Mayer’s Hematoxylin Solution (FUJIFILM Wako Pure Chemical Corporation) at RT for 3 min and in Eosin Alcohol solution (FUJIFILM Wako Pure Chemical Corporation) at RT for 1 min. Dehydration was performed using graded ethanol solutions (70%, 80%, 90%, and 100%) and clearing was performed three times using xylene.

### Cochlea whole-mount preparation

The inner ears were dissected from mice heads at E18.5 and fixed in 4% PFA in PBS at RT for 1h. After fixation and before primary antibody staining, the outer membrane, including the tectorial membrane, was removed to expose the organs of Corti. After staining with a secondary antibody, the organ of Corti was dissected and mounted on a glass slide for imaging.

### Proliferation and apoptosis assays

Cell proliferation in the cochlea was assessed by detecting the incorporated EdU (Thermo Fisher Scientific) on frozen sections. EdU was detected using the Click-iT Plus EdU Cell Proliferation Kit for Imaging Alexa 555 Dye (Thermo Fisher Scientific), according to the manufacturer’s instructions. Pregnant mice were injected with EdU (Thermo Fisher Scientific) at E12.5, E13.5, and E14.5 (three injections at 50 μg/g at 2-h intervals) and at E16.5 (a single injection at 100 μg/g). E12.5, E13.5, and E14.5 embryos were collected 8 h after the first injection. E16.5 embryos were collected 4 h after the injection. The basal or basal-to-middle regions of the cochlear duct were observed at E12.5 or at E13.5, E14.5, and E16.5, respectively.

Apoptotic cells were detected by identifying the expression of CC3 in frozen sections via IHC staining.

### ABR and DPOAE

ABR measurements were performed under general anesthesia as described previously (Kada et al., 2009) at P21 (n = 3 for each genotype). The thresholds for 10, 20, and 40 kHz were determined based on the responses at different intensities with 5 dB sound pressure level intervals. DPOAE recordings were performed as described previously (Hamaguchi et al., 2012) at P21 (n = 4 for each genotype). Two primary tones (f1, f2, f1<f2) were used as input signals, with f2 set at eight frequency points (4, 6, 8, 12, 16, 24, 32, and 40 kHz), maintaining a frequency ratio of f2/f1 = 1.2. The intensity levels of the stimulatory sounds were 65 and 55 dB sound pressure level for f1 and f2, respectively. DPOAE was detected as a peak at 2f1–f2 in the spectrum.

### Quantification

Cell quantification and measurements were performed at E18.5 using the Cell Counter plugin of ImageJ (Schneider et al., 2012). The total length of the cochlea was measured based on the region with MYO7A-positive hair cells from the basal to apical turns. Cochlear hair cells were identified by phalloidin and MYO7A labeling. Two types of cells, PROX1- and SOX2-positive cells, were counted to quantify the supporting cells of the cochlea. The cochlear duct was divided into three regions: basal, middle, and apical, and we counted the number of cells per 200 µm in each region for MYO7A- and PROX1-positive cells or 100 µm in the basal region for SOX2-positive cells.

### Statistical analysis

For all statistical analyses, at least three samples from each experimental group were analyzed.

Student’s *t*-test was used to determine the differences between two experimental groups. One-way or two-way analysis of variance was performed to assess the differences between more than two experimental groups and *p*-values less than 0.05 were considered statistically significant. Statistical analyses were performed using R version 4.2.2 (2022-10-31). All details of statistical analyses are provided in the figures and legends.

## Acknowledgements

We thank Dr. Rudolf Grosschedl for providing Ebf1^-/-^ and Ebf1^fl/fl^ mice and Dr. Ryoichiro Kageyama for providing *Atoh1* ISH probe. This study was supported by JSPS KAKENHI (grant number: JP22H03234) awarded to NY.

## Author Contributions

N.Y. and H.O. designed the study. H.K., H.O., R.Y., and Y.T. performed the experiments with advice from N.Y., T.K., and K.O.. A.Y. performed ABR and DPOAE. H.K. and N.Y. wrote the manuscript with contribution from all authors.

## Supplemental Figures and Legends for Supplemental Figures

**Supplemental Fig. S1.**
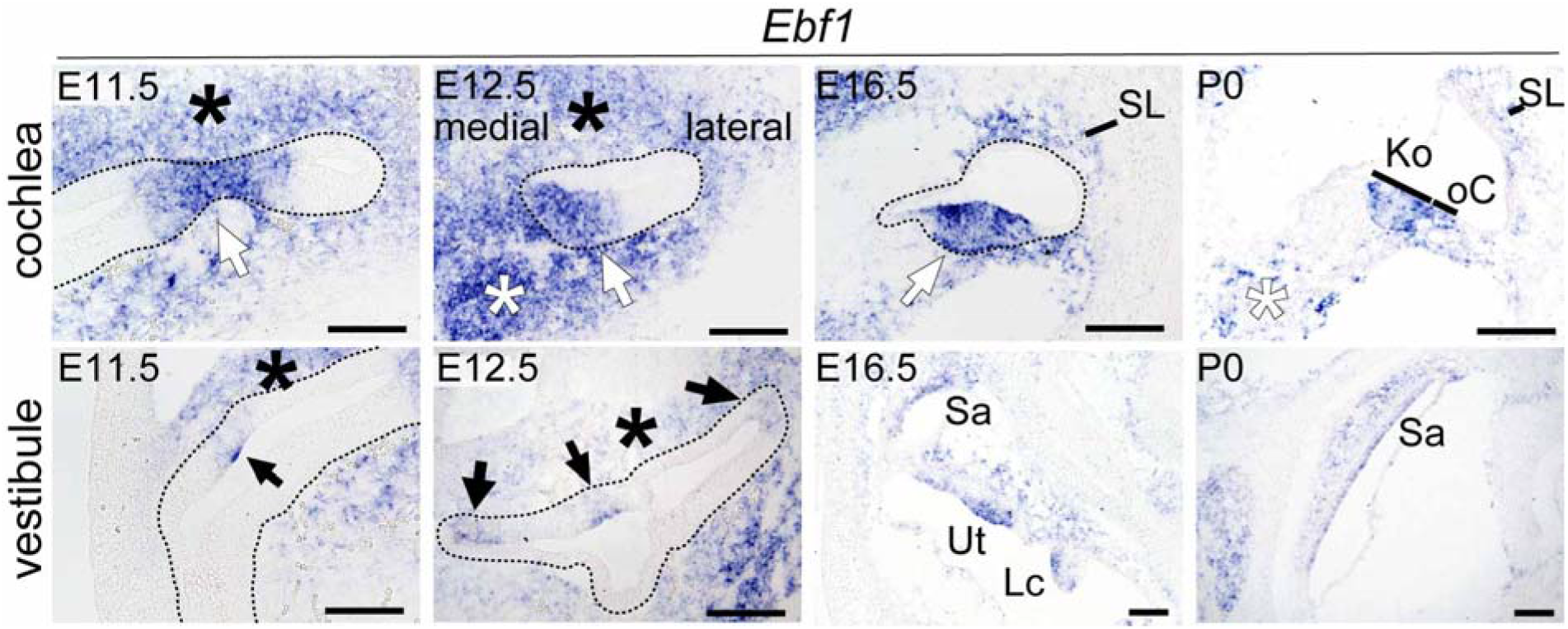
Spatiotemporal expression of *Ebf1* during inner ear development. Result of *in situ* hybridization for *Ebf1* and *Sox2* on cross sections of the inner ear of wild-type mice at embryonic day (E) 11.5, E12.5, E16.5, and P0. Areas enclosed by dashed lines indicate the inner ear epithelium. From E11.5 to P0, *Ebf1* is expressed in the sensory epithelium of the cochlea (white arrows), the vestibular and semicircular canals (arrows), the spiral ganglion (white asterisks), and the surrounding mesenchymal tissues (asterisks and SL). sg, spiral ganglion; Ut, utricle; Sa, saccule; Lc, lateral crista; SL, spiral ligament; Ko, Kölliker’s organ; oC, organ of Corti. Scale bars: 100 μm.

**Supplemental Fig. S2.**
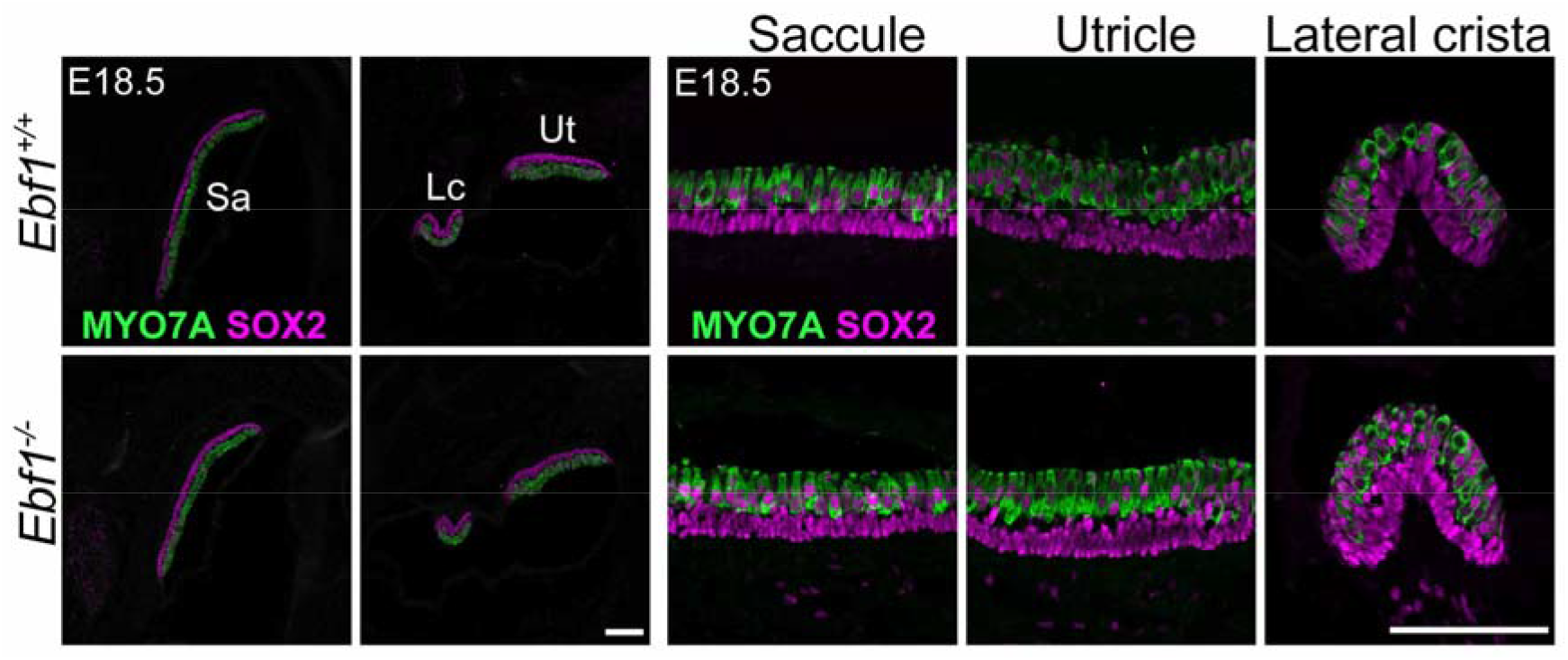
*Ebf1* deletion does not affect hair and supporting cells in the vestibular macula and crista. Cross sections of the embryonic day (E) 18.5 vestibular region immunostained for MYO7A (green) and SOX2 (magenta). Low-magnification (left four panels) and high-magnification (right panels) images of the saccule, lateral crista, and utricle are presented. Ut, utricle; Sa, saccule; Lc, lateral crista. Scale bars: 100 μm.

**Supplemental Fig. S3.**
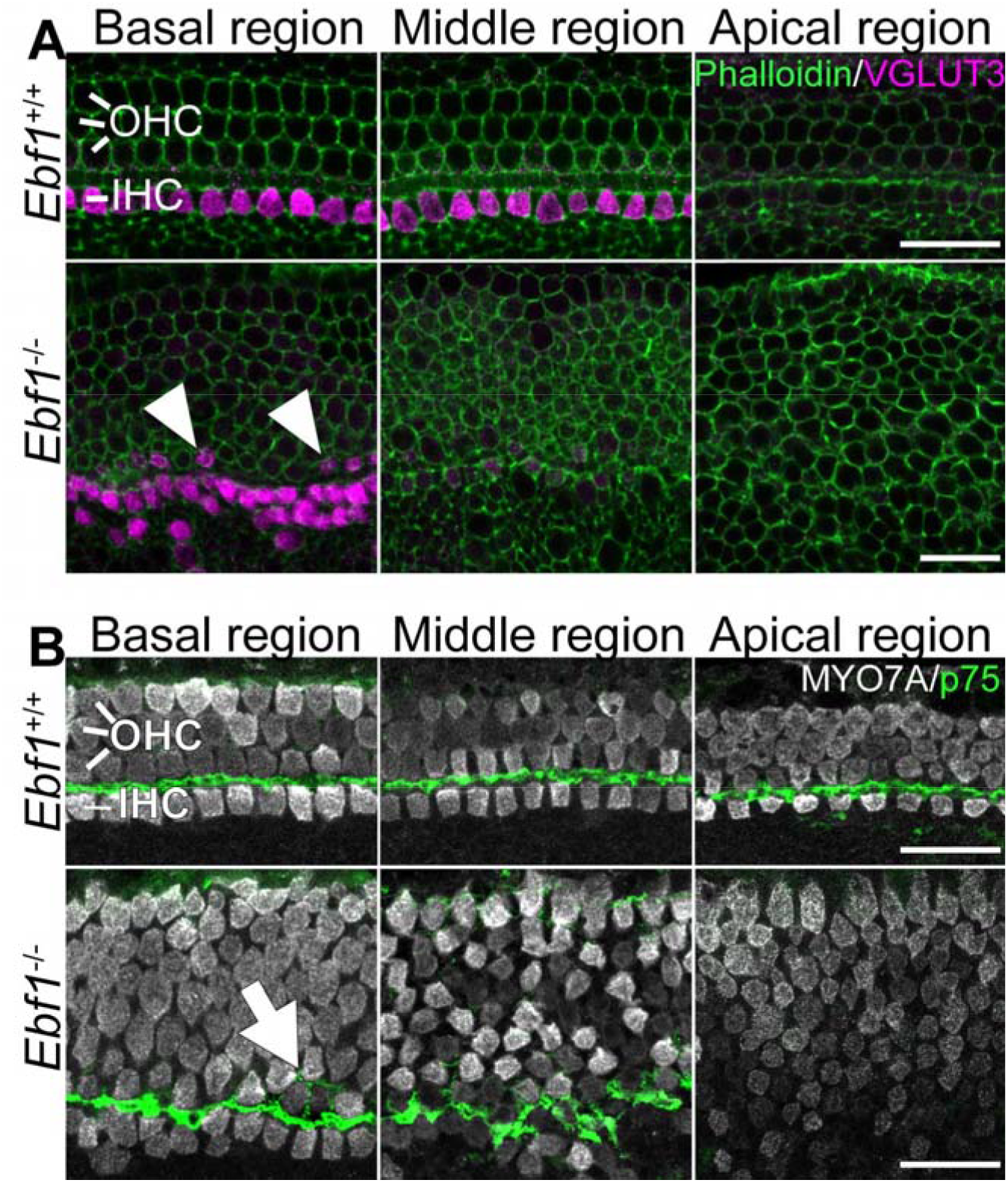
*Ebf1* deletion causes increased number of inner hair cells and delayed differentiation of hair and supporting cells. High-magnification images of the basal, middle, and apical regions of whole-mount cochlear samples from E18.5 *Ebf1^+/+^* and *Ebf1^-/-^* mice. **A**. Immunostaining with phalloidin (green) and an anti-VGLUT3 antibody (magenta). In the cochlea of *Ebf1^-/-^* mice, the row number of VGLUT3-positive cells was 2 to 3, whereas this number was only 1 in wild-type mice. VGLUT3-positive cells were observed only in the basal region of the cochlea but not in the middle and apical regions of *Ebf1^-/-^* mice. Several VGLUT3-positive cells were observed in the outer hair cell area (arrowheads). **B**. Immunostaining with anti-MYO7A (gray) and p75 (green) antibodies. In the cochlea of *Ebf1^-/-^* mice, the row number of p75-positive cells was similar to that in *Ebf1^+/+^*mice, although their arrangement was deteriorated in *Ebf1^-/-^* mice (arrow). p75-positive cells are observed only in the basal and middle region in the *Ebf1^-/-^* mice. Scale bars: 20 μm.

**Supplemental Fig. S4.**
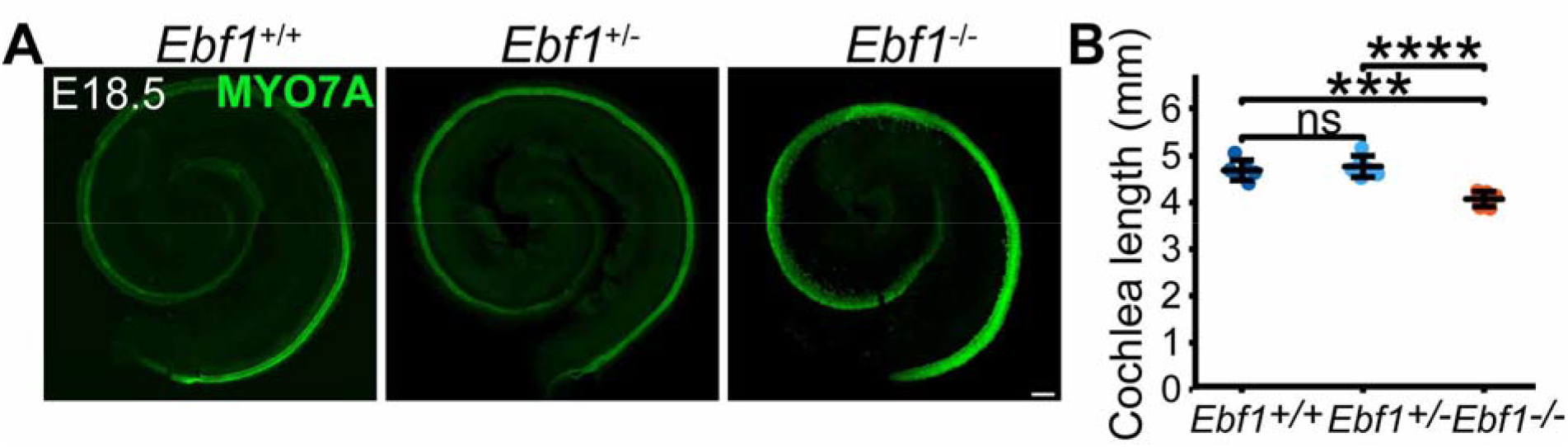
Quantification of cochlear length. **A**. Whole mount images of the cochlea of embryonic day (E) 18.5 *Ebf1^+/+^*, *Ebf1^+/-^*, and *Ebf1^-/-^* mice labeled with MYO7A (green). **B**. Quantification of cochlear duct length of E18.5 *Ebf1^+/+^*, *Ebf1^+/-^*, and *Ebf1^-/-^* mice. The cochlear length of *Ebf1^-/-^* mice was significantly shorter than those of *Ebf1^+/+^*and *Ebf1^+/-^*mice. Two-way analysis of variance (ANOVA) test with Bonferroni *post-hoc* tests was performed. \*\*\**p* < 0.001, and \*\*\*\**p* < 0.0001; ns, not significant. Error bars represent mean ± standard deviation. n = 6

**Supplemental Fig. S5.**
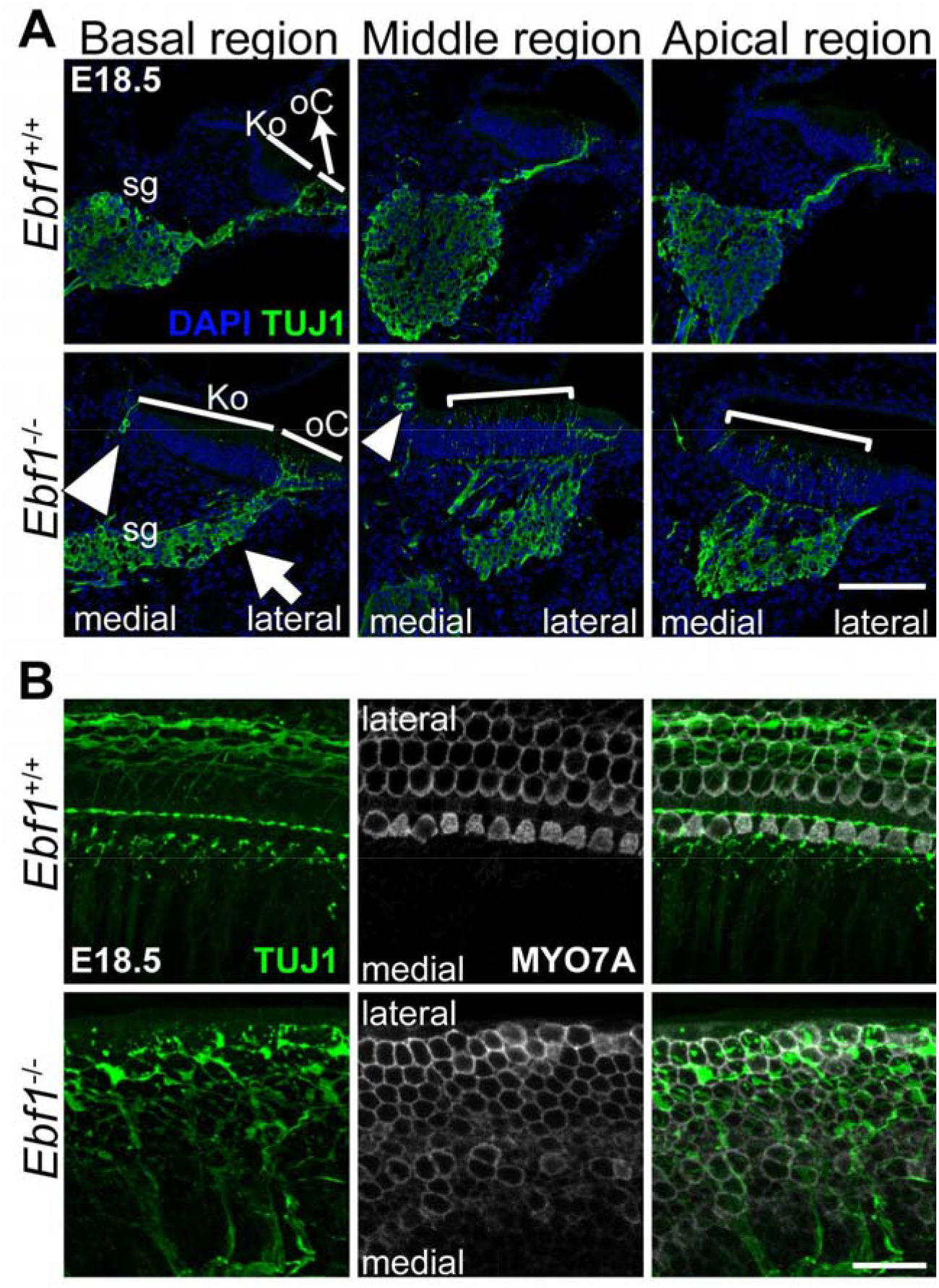
*Ebf1* deletion causes aberrant spiral ganglion development and axon outgrowth to cochlear hair cells. **A**. Cross sections of the basal, middle, and apical regions of the cochlea and spiral ganglion in embryonic day (E) 18.5 *Ebf1^+/+^* and *Ebf1^-/-^* mice stained with Tubulin β 3 (TUJ1, green) and 4’,6-diamidino-2-phenylindole (DAPI, gray). The morphology of the spiral ganglion in *Ebf1^-/-^* mice differed from that in *Ebf1^+/+^* mice. TUJ1-positive cell bodies were observed below the organ of Corti (arrow) and at their normal site. Additionally, innervation from the spiral ganglion was observed in the hair cell part and in Kölliker’s organ in the middle and apical of the cochlea (arrowheads and brackets). **B**. High-magnification view of the basal region of the whole mount cochlear image in E18.5 *Ebf1^+/+^* and *Ebf1^-/-^* mice labeled with TUJ1 (green) and MYO7A (gray). In *Ebf1^+/+^* mice, the neurons ran parallel to the outer hair cells, whereas in *Ebf1^-/-^* mice, the neurons formed a reticulation within the cochlear hair cell regions. Scale bars: 100 μm (**A**) and 20 μm **(B**).

**Supplemental Fig. S6.**
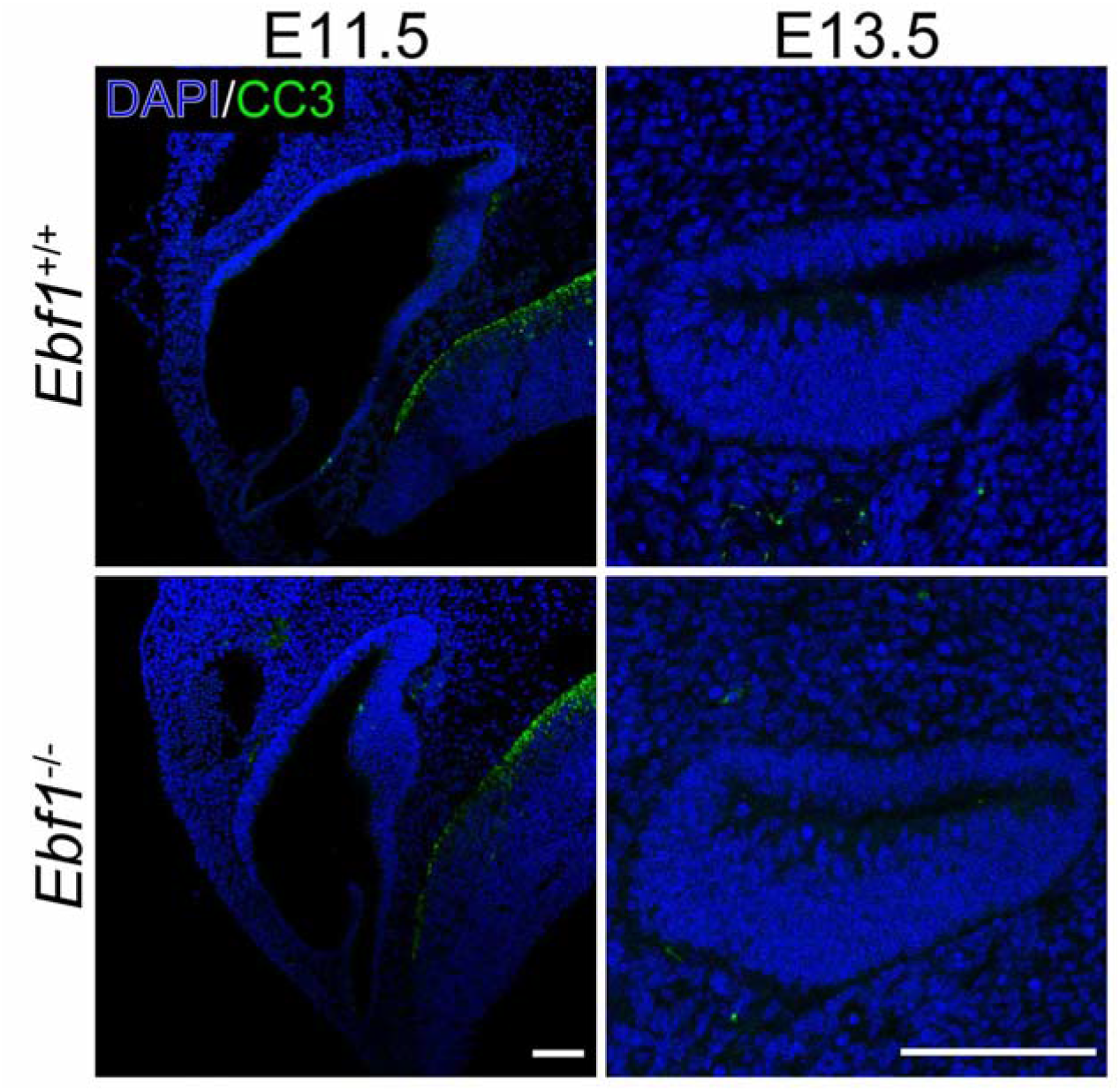
*Ebf1* deletion did not affect apoptosis during inner ear development. Cross sections of the embryonic day (E) 11.5 inner ear and E13.5 cochlear basal region from *Ebf1^+/+^*and *Ebf1^-/-^* mice. Sections were labeled with cleaved caspase 3 (CC3, green) and 4’,6-diamidino-2-phenylindole (DAPI, blue). The number of CC3-positive cells was similar between *Ebf1^-/-^* and *Ebf1^+/+^*mice at E11.5 and E13.5. Scale bars: 100 μm.

